# Probing Bioactive Chemical Space to Discover RNA-Targeted Small Molecules

**DOI:** 10.1101/2023.07.31.551350

**Authors:** Sarah L. Wicks, Brittany S. Morgan, Alexander W. Wilson, Amanda E. Hargrove

**Affiliations:** Department of Chemistry; Duke University; 124 Science Drive; Durham, NC 27708; Department of Chemistry & Biochemistry; University of Notre Dame; 123 McCourtney Hall Notre Dame, IN 46556

**Author notes:** Co-first authors (Sarah L. Wicks, Brittany S. Morgan).

## Abstract

Small molecules have become increasingly recognized as invaluable tools to study RNA structure and function and to develop RNA-targeted therapeutics. To rationally design RNA-targeting ligands, a comprehensive understanding and explicit testing of small molecule properties that govern molecular recognition is crucial. To date, most studies have primarily evaluated properties of small molecules that bind RNA *in vitro*, with little to no assessment of properties that are distinct to selective and bioactive RNA-targeted ligands. Therefore, we curated an RNA-focused library, termed the <u>D</u>uke <u>R</u>NA-<u>T</u>argeted <u>L</u>ibrary (DRTL), that was biased towards the physicochemical and structural properties of biologically active and non-ribosomal RNA-targeted small molecules. The DRTL represents one of the largest academic RNA-focused small molecule libraries curated to date with more than 800 small molecules. These ligands were selected using computational approaches that measure similarity to known bioactive RNA ligands and that diversify the molecules within this space. We evaluated DRTL binding *in vitro* to a panel of four RNAs using two optimized fluorescent indicator displacement assays, and we successfully identified multiple small molecule hits, including several novel scaffolds for RNA. The DRTL has and will continue to provide insights into biologically relevant RNA chemical space, such as the identification of additional RNA-privileged scaffolds and validation of RNA-privileged molecular features. Future DRTL screening will focus on expanding both the targets and assays used, and we welcome collaboration from the scientific community. We envision that the DRTL will be a valuable resource for the discovery of RNA-targeted chemical probes and therapeutic leads.

## INTRODUCTION

The use of small molecules to modulate RNA structure and function has become invaluable for studying the many roles of RNA in biological processes^1^ as well as discovering effective therapeutic treatments.^2-5^ Early on, the promise of RNA as a ‘druggable’ target was demonstrated by the identification of several RNA-targeting small molecule classes such as aminoglycosides.^6, 7^ These ligands bound highly abundant and structured ribosomal RNA (rRNA) and inhibited protein synthesis in prokaryotes, leading to the development of numerous FDA-approved antibiotics.^8^ Over the last 25 years, momentous efforts to target non-ribosomal RNAs, which are less studied and have lower abundance (< 10% of total RNA), have yielded a modest collection of ∼150 biologically active small molecules that target bacterial, fungal, viral, and mammalian RNAs.^9, 10^ Notable examples include Phase 2 clinical trial candidate Branaplam^11^ and recently FDA-approved Risdiplam^12^, both of which were developed to target mRNA splicing and treat spinal muscular atrophy. While immensely promising, the field of targeting RNA with small molecules is still in its infancy; we have only begun to elucidate and evaluate the guiding principles that govern RNA:small molecule recognition.

Collective efforts to identify the properties of small molecules that selectively interact with RNA have yielded critical insights into molecular recognition. For example, the Disney laboratory first utilized selection-based, two-dimensional combinatorial screens to identify chemotypes that bind RNA with high affinity and selectivity.^13^ Enriching libraries with these RNA-binding chemotypes has led to increased hit rates and the identification of bioactive small molecules for RNA.^14-16^ Recently, large RNA-targeted screens by academic and industrial laboratories revealed differentiating properties between *in vitro* RNA-binding small molecules and compounds that were known to bind proteins^17^ and/or explicitly known to not bind RNA.^18^,^19^ These works largely corroborated our analysis from 2017, where we examined a collection of small molecules in the <u>R</u>NA-targeted <u>BI</u>oactive liga<u>N</u>d <u>D</u>atabase, termed R-BIND (SM) (n = 67). We identified cheminformatic descriptors related to molecular structure, complexity, and recognition that distinguished the biologically active, non-ribosomal RNA-targeted small molecules from FDA-approved drugs, which largely (∼90%) target proteins.^20^ Similar to FDA-approved drugs, the majority of R-BIND (SM) complied with medicinal chemistry rules for drug-likeness; however, biologically active RNA-targeted small molecules had distinctive structural features, including an increased nitrogen count, decreased oxygen count, decreased flexibility, and increased counts of aromatic and heteroatom-containing rings.^21^ Notably, while the size of R-BIND (SM) increased by ∼225% (n = 153) over three years, little change in the distributions of the cheminformatic descriptors was observed, further supporting the existence of a privileged chemical space for bioactive RNA-targeted small molecules.^9, 10^

One approach to further examine RNA-biased small molecule chemical space is to develop and evaluate RNA-focused libraries. Indeed, about 40% of R-BIND (SM) ligands were discovered using a focused screening approach, often yielding higher hit rates as compared to high-throughput screens.^9, 10, 22^ Design strategies for focused libraries have included enrichment with scaffolds known to bind RNA^14, 15, 23, 24^ as well as fragmentation of^25^ or chemical similarity to^17, 18, 26^ known RNA-binding small molecules. While successful, these approaches are currently limited to building libraries based on small molecules known to bind RNA *in vitro*. As a result of our continuous efforts to curate and analyze R-BIND (SM),^9, 10, 21, 22^ we were uniquely positioned to develop and evaluate an RNA-focused library based upon molecular properties of bioactive, RNA-targeted small molecules. The inclusion of features that are prominent in biologically active small molecules was proposed to increase selectivity and expedite the development of chemical probes for RNA.

Herein, we describe the curation and evaluation of an RNA-focused library biased towards the physicochemical and structural properties of biologically active, RNA-targeted small molecules. Using 20 cheminformatic parameters, a k-Nearest Neighbor algorithm was applied to examine and select commercial small molecules that had similar features to R-BIND (SM) ligands. During the selection process, we discovered that particular regions of R-BIND (SM) chemical space were either inaccessible or sparsely occupied by commercial libraries. Purchased small molecules, which we termed the <u>D</u>uke <u>R</u>NA-<u>T</u>argeted <u>L</u>ibrary (DRTL), were evaluated against a panel of four RNAs, utilizing two optimized fluorescent indicator displacement assays. By leveraging R-BIND (SM), the DRTL led to hit rates similar to RNA-focused screens^9, 10, 22^ without prior knowledge of a particular RNA-binding ligand or scaffold. Further, we successfully identified multiple RNA-binding small molecules to pursue for structure-activity relationship studies and biological assays. Additional screens will provide further insights into biologically-relevant RNA chemical space, enable explicit testing of the guiding principles we previously identified,^9, 10, 21^ and contribute to the continued identification of RNA-privileged scaffolds and molecular features, all of which will facilitate the imperative discovery of chemical probes for RNA.

## METHODS

Details of materials, methods, and experimental procedures are provided in the Supporting Information.

## RESULTS AND DISCUSSION

### Curation of an RNA-focused Library

To identify commercial small molecules that have similar molecular properties to R-BIND (SM), we utilized a k-Nearest Neighbor (k-NN) algorithm.^9, 27^ We defined the chemical space of R-BIND (SM) v1.1 (n = 75) by a set of 20 cheminformatic parameters that was previously used to analyze R-BIND (SM) ligands and compare them to: i) RNA binders without reported bioactivity and ii) FDA-approved drugs, a surrogate for protein-targeted, bioactive ligands.^21^ The parameters included medicinal chemistry, structural complexity, and molecular recognition descriptors (SI Table 1). The R-BIND (SM) ligands were plotted in the 20-dimensional chemical space and the Euclidean distance between every R-BIND (SM) pair was calculated. For each R-BIND (SM) ligand, the shortest distance to its nearest neighbor was identified. The distance ranged from 0.3418 to 7.3373 with an average shortest distance of 2.1923 (SD Table 1). Notably, there were regions within the chemical space that had multiple R-BIND (SM) ligands in proximity, with 25/75 ligands having at least two nearest neighbors within the average distance (defined henceforth as NNs) and of these, one R-BIND (SM) ligand had seven NNs (SD Table 2).

Commercially available small molecules from ChemBridge and ChemDiv libraries (n = 2,484,294) were plotted in the 20-dimensional chemical space defined by R-BIND (SM) and evaluated for R-BIND-like character (Figure 1A). The k-NN algorithm was applied using the R-BIND-like distance, defined as the average shortest distance (d = 2.1923), to identify commercial ligands that occupied similar chemical space.^9^ In total, we identified 1,096,020 NN pairs between a subset of commercial small molecules (n = 572,145) and R-BIND (SM) ligands (n = 61) (SD Table 3). The number of NNs for these R-BIND (SM) ligands ranged from 1 to 197,433 small molecules with a median of 1,057 (SI Table 2). Notably, for 14 of the 75 R-BIND (SM) ligands, there were no commercial small molecules nearby in chemical space (SI Table 3). The majority of these (n = 9) were also isolated from other R-BIND (SM) ligands in chemical space (d > 2.1923) (SI Table 4). Altogether, our analysis indicates that while commercially available R-BIND-like small molecules can be found, there are regions of R-BIND (SM) chemical space that are sparsely populated, if at all, by commercial ligands.

**Figure 1.**
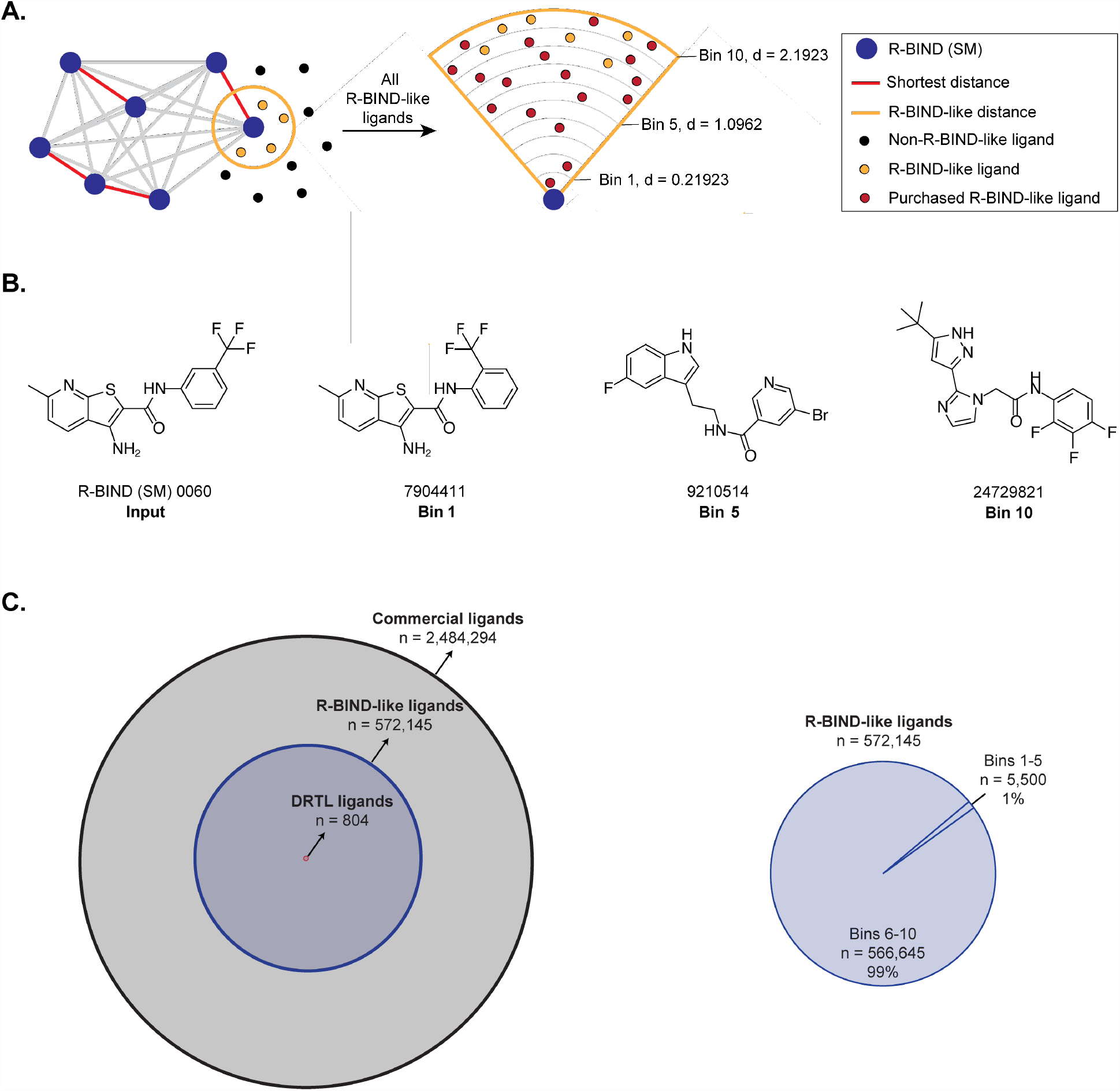
Curation of the Duke RNA-Targeted Library (DRTL). (A) Schematic of the curation of the DRTL. The R-BIND (SM) library is plotted in 20-dimensional chemical space defined by cheminformatic parameters. Using a k-Nearest Neighbor algorithm, the Euclidean distance of each R-BIND (SM) to all of its neighbors is calculated. Then, the shortest nearest neighbor distance for each R-BIND (SM) is averaged. This R-BIND-like distance is used to identify commercial ligands that occupy similar chemical space. All R-BIND-like ligands for a given R-BIND (SM) were grouped into one of ten bins, according to their nearest neighbor distance. Up to three ligands from each bin were randomly selected for purchase. (B) Example of a nearest neighbor search. Commercial ligands that were within the average distance (d = 2.1923) to R-BIND (SM) 0060 are shown and represent selections from Bins 1, 5, and 10. (C) Summary of total commercial ligands that were evaluated for R-BIND-likeness (n = 2,484,294), found to be R-BIND-like (n = 572,145), and purchased for the DRTL (n = 804). Pie graph (right) represents the percentage of commercial R-BIND-like ligands that resided either in one of the innermost (Bins 1-5, d = 0.0000-1.0962) or outermost (Bins 6-10, d = 1.0963-2.1923) bins to R-BIND (SM).

To further assess the similarity to R-BIND (SM), commercially available R-BIND-like small molecules were sorted by measured distances into one of ten bins, where Bin 1 was shortest in distance (d < 0.2192) and Bin 10 was farthest in distance (d = 1.9732-2.1923) to an R-BIND small molecule. The distance of a commercial ligand to an R-BIND (SM) is proportional to its similarity in terms of the 20 cheminformatic parameters (Figure 1B). The majority of NNs (n = 1,090,213, 99.5%) resided in the outermost bins (Bins 6-10, d = 1.0963-2.1923) (Figure 1C, SD Table 4). The innermost bins (Bins 1-5, d = 0.0000-1.0962) consisted of only 5,807 NN pairs between a fraction of commercial ligands (n = 5,500) and a subset of R-BIND (SM) (n = 30). Notably, the lack of commercial compounds in the innermost bins limits the assessment of small molecules that most closely resemble the molecular properties of R-BIND (SM) ligands.

Commercially available small molecules that represented bioactive RNA-targeted chemical space were then purchased to build the <u>D</u>uke <u>R</u>NA-<u>T</u>argeted <u>L</u>ibrary (DRTL). Up to three commercial compounds were randomly selected in each of the 10 bins for every R-BIND (SM) ligand (n = 75). While a commercial compound could be a NN to multiple R-BIND small molecules, the compound was selected only once to maximize library diversity and size. Utilizing this method, a maximum of 2,250 commercial small molecules could have been selected. However, NNs were identified for only a subset of R-BIND (SM) ligands (n = 61) and the majority of bins had less than three unique ligands (SD Tables 5 and 6). In total, only one-third of the maximum number of ligands could be selected and purchased as the DRTL (n = 804) (Figure 1C and SD Table 7). These results further illustrate the need to supplement libraries with novel synthetic analogs that expand the chemical space coverage and density, especially in regions nearest to R-BIND (SM) ligands.

### Physicochemical, Spatial, and Structural Analysis of the DRTL

To compare the DRTL and R-BIND (SM), the distributions of the 20 cheminformatic parameters were analyzed using a Mann-Whitney U statistical test. As expected, the DRTL and R-BIND (SM) had similar distributions for the majority of the cheminformatic parameters. Statistically significant differences were observed for eight of the twenty descriptors (SI Table 5); however, the exclusion of R-BIND (SM) ligands that had no commercial NNs (n = 14) resulted in statistically significant differences for only five of the twenty parameters (SI Table 6). These differences included a reduction in number of hydrogen bond donors (HBD), ring complexity (SysRR), and total charge (TC) for the DRTL. Furthermore, decreased water solubility (higher log *P*, log *D*) was observed for the DRTL. The differences found between the libraries (DRTL and RBIND) is likely due in part to the lack of commercially available ligands in close proximity to many R-BIND (SM) ligands.. Using principal component analysis,^28^ the regions of R-BIND (SM) chemical space that were encompassed by the DRTL were visualized (Figure 2A). As anticipated, most of R-BIND (SM) chemical space was populated by the DRTL, though with varying coverage of density and proximity. Thus, the DRTL possesses R-BIND-like character and is representative of a subset of the biologically active, RNA-targeted chemical space defined by R-BIND (SM).

**Figure 2.**
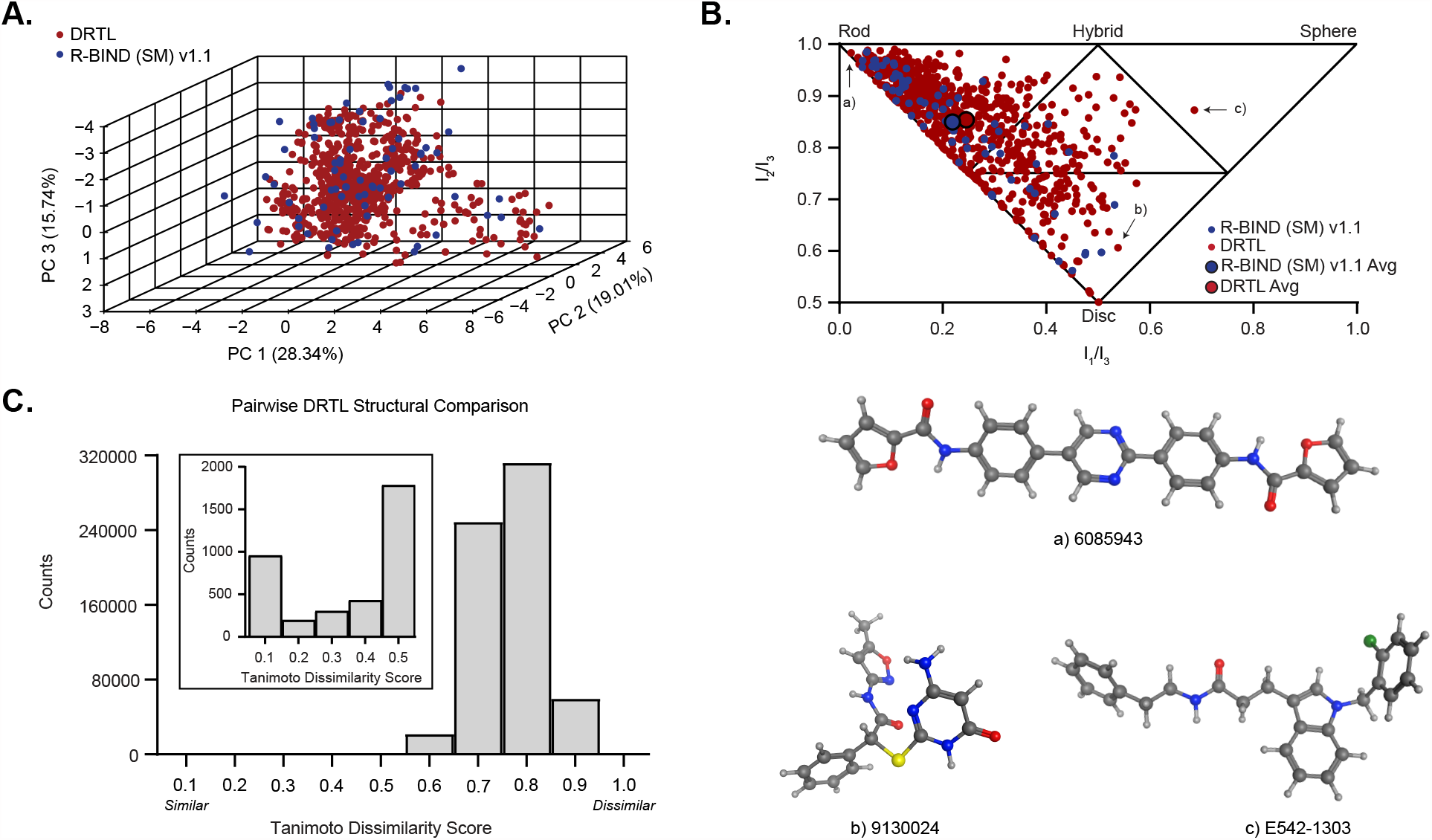
Cheminformatic analysis of the DRTL. (A) Principal component analysis plot of principal components (PC) 1-3 based on 20 cheminformatic parameters calculated for R-BIND (SM) v1.1. The DRTL was added as supplemental data and does not contribute to PCs. The principal component and subsequent percent contribution is indicated on each axis. (B) Normalized principal moments of inertia ratios for DRTL and R-BIND (SM). Each point represents the Boltzmann average of a molecule using conformations within 3 kcal/mol of the lowest energy conformer. Larger circles outlined in black represent the library averages. Structures of DRTL ligands that represent a) rod-like (6085943), b) disc-like (9130024), and c) sphere-like (E542-1303) shapes are shown. (C) Distribution of Tanimoto dissimilarity scores of small molecules in the DRTL. Tanimoto dissimilarity scores from pairwise comparisons of the 804-member library were used to create the histogram. A score of 1.0 indicates that two ligands are highly dissimilar. The inset shows the lower counts for Tanimoto dissimilarity scores of 0.1-0.5.

Additionally, the spatial properties of the DRTL were evaluated and compared to R-BIND (SM). The 3-dimensional (3D) shape of each small molecule was calculated using principal moments of inertia (Figure 2B).^29^ The library averages for the DRTL and R-BIND (SM) were nearly identical, and partitioning of the libraries into four sub-triangles showed similarities in the library percentages that populated the rod (∼75%), disc (∼15%), sphere (< 1%), and hybrid (∼10%) triangles (SI Tables 8 and 9). To quantitatively compare the library distributions, the Kolmogorov-Smirnov (KS) statistical test was used. The KS test revealed that the shape distributions of the DRTL are statistically different from R-BIND (SM), containing more disc- and sphere-like and less rod-like character (SI Table 10). These observations are supported by cumulative distance distribution graphs, which measure the distance of each small molecule from the rod (0,1), disc (0.5,0.5), or sphere (1,1) vertices (SI Figure 1). Similar results were obtained when R-BIND (SM) ligands that had no commercial NNs were excluded from the analysis (SI Table 11). Due to the computational resources required to calculate predicted small molecule conformations for all NN pairs, 3D shape was not considered in the curation of the DRTL yet our analysis indicates that the DRTL does contain compounds with shapes similar to bioactive RNA ligands.

To evaluate the structural diversity of DRTL, chemical hashed fingerprints that describe molecular structure were calculated. Small molecule fingerprints were compared to one another using a Tanimoto dissimilarity score, where a score of 1 indicates that two ligands are highly dissimilar.^30^

For DRTL, the average Tanimoto dissimilarity score for all pairwise comparisons was 0.76 ± 0.03, suggesting that library members are generally chemically diverse (Figure 2C).^31^ However, ∼25% of the DRTL (n = 221/804) had a low Tanimoto dissimilarity score (≤ 0.30) for at least one pairwise comparison, indicating that common scaffolds are found among some of the library members. Comparisons between the DRTL and R-BIND (SM) revealed that the two libraries are structurally dissimilar, with less than 5% of the DRTL (n = 30/804) having Tanimoto dissimilarity scores ≤ 0.30 for pairwise comparisons with R-BIND (SI Figure 2). This analysis confirmed that the DRTL is comprised of novel chemical matter and has the potential to identify RNA-targeting small molecules with distinct molecular structures.

### *In Vitro* High-throughput Screening of the DRTL

As a preliminary evaluation of our library selection strategy, we investigated *in vitro* binding of the DRTL to four biologically-relevant RNAs: truncated constructs of the transactivation response element (TAR) in type 1 and 2 of the human immunodeficiency virus (HIV)^32^ as well as rCAG and rCUG expansions that contain repeat sequences associated with Huntington’s disease and Myotonic Dystrophy Type 1,^33^ respectively. The HIV TAR RNAs adopt a stem-loop secondary structure that includes a 6-nucleotide (nt) hairpin loop with either a 2-nt (HIV-2 TAR) or 3-nt (HIV-1 TAR) bulge (Figure 3A). The expanded repeats form five 1 x 1-nt internal loops comprised of either AA (rCAG_12_) or UU (rCUG_12_) sequences with 4-nt closing hairpin loops. Notably, approximately 30% of R-BIND (SM) ligands target either HIV-1 TAR (n = 14) or CUG expanded repeats (n = 8). These R-BIND (SM) ligands also constituted NN pairs with a substantial subset of the DRTL (n = 446, 55%) (SD Table 8). While CAG expanded repeats had only one reported R-BIND (SM) ligand and HIV-2 TAR had none, their structural similarity to CUG expanded repeats and HIV-1 TAR, respectively, could provide valuable insights into ligand selectivity as well as relevant small molecule chemical space for targeting these RNA structures. Collectively, the four RNA targets selected are ideal for initial screens of the DRTL with the goal of generating binding and selectivity profiles.

**Figure 3.**
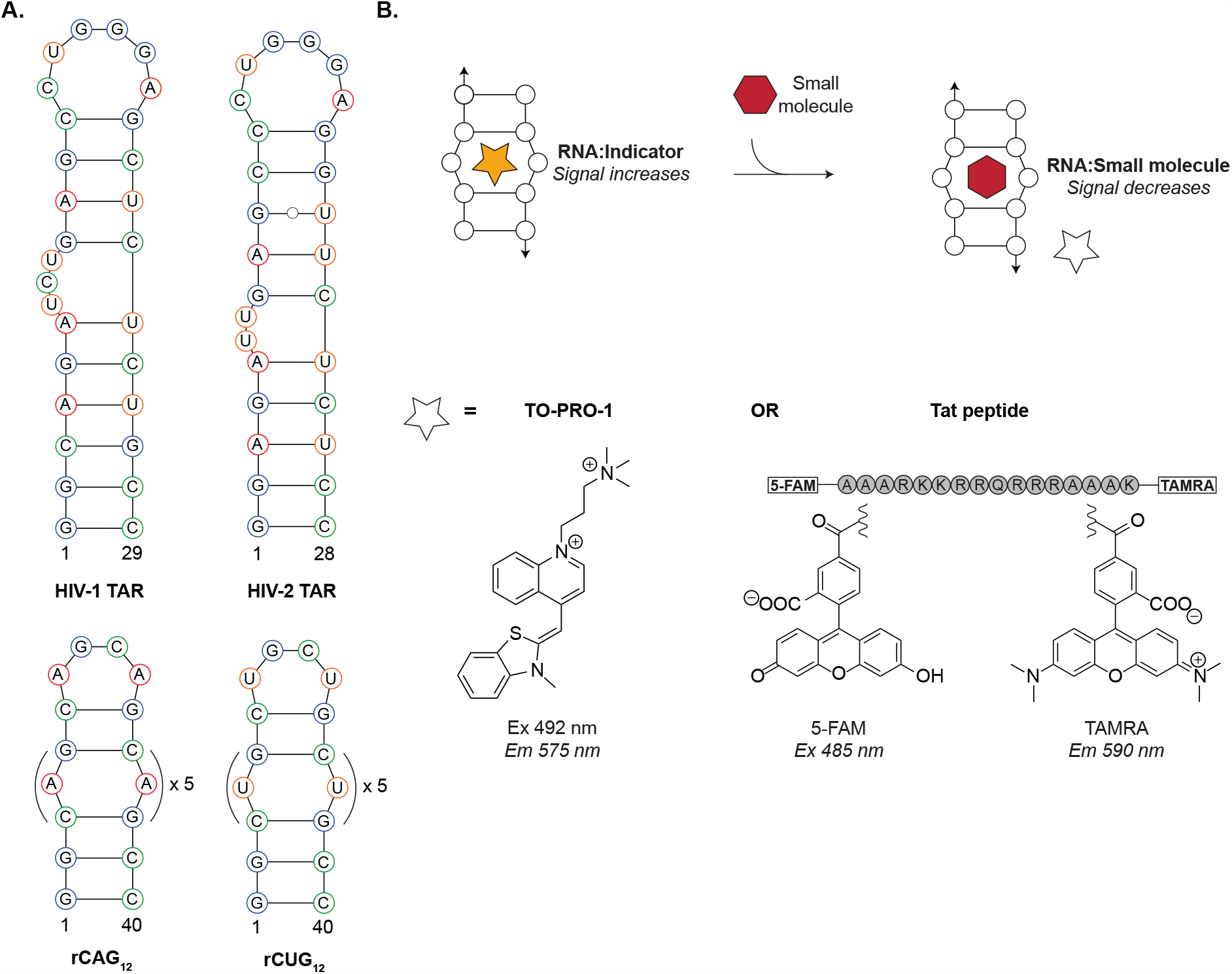
Selection of RNA targets and primary screening assays. (A) Sequences and secondary structures of RNA targets. (B) TO-PRO-1 and Tat peptide indicator displacement assays. When the indicator is bound by RNA, signal is enhanced. Displacement of the indicator by a competitive small molecule diminishes signal. TO-PRO-1 was used to screen the DRTL against all four RNA targets. Tat peptide, labeled with 5-FAM at the N-termini and TAMRA at the C-termini, was used to screen the DRTL against HIV-1 and HIV-2 TAR RNA targets. Structures of TO-PRO-1, 5-FAM and TAMRA fluorophores with respective excitation (Ex) and emission (Em) wavelengths as well as sequence of Tat peptide are shown. Amino acid abbreviations: A = Alanine, R = Arginine, K = Lysine, Q = Glutamine.

For the screening approach, we selected two different fluorescence-based indicator displacement assays (IDAs) (Figure 3B). The first assay uses the fluorescent indicator TO-PRO-1, an intercalating cyanine dye that increases fluorescence upon binding nucleic acids.^34^ TO-PRO-1 has been successfully used in screens against RNAs such as HIV-1 TAR and repeat expansions and thus, was employed against all four RNA targets.^14, 35-38^ The second assay utilizes an arginine rich, truncated peptide of Tat protein that binds primarily by hydrogen bonding and electrostatic interactions to the phosphate backbone of RNAs.^39, 40^ The Tat peptide is labeled with a Förster Resonance Enhancement Transfer pair, namely 5-carboxyfluorescein donor and 5-carboxytetramethylrhodamine acceptor fluorophores, for which signal intensity increases when the peptide is bound to RNA.^41^ The Tat peptide construct has been used as a probe to identify RNA-binding ligands mainly for HIV RNAs^38, 41-43^ and therefore, was employed against HIV-1 and HIV-2 TAR targets only. By evaluating the DRTL with two different IDAs, we anticipated that our results would reveal potential screening bias and increase confidence in hits that are identified.

Both IDAs were optimized for the selected RNA targets and assessed for high-throughput assay quality. The assays were conducted in a physiologically relevant buffer containing potassium (140 mM), sodium (10 mM), and magnesium (1 mM)^44^ chloride salts as well as 0.01% Triton-x-100, 5% DMSO, and 0.1 mM EDTA (pH = 7.2). Under these conditions, apparent dissociation constants of the indicators to RNA targets were low micromolar for TO-PRO-1 (0.71-1.18 μM) and low nanomolar for Tat peptide (92-188 nM) (SI Tables 14 and 15, SI Figures 3 and 4). Next, the quality of the IDAs was quantified by a Z’-Factor, a measure that reflects both the signal dynamic range and variation of measurements.^45^ The initial fraction of bound (Fb^0^) indicator can influence the signal dynamic range and sensitivity of an IDA.^46, 47^ With TO-PRO-1, an RNA concentration that corresponded to an Fb^0^ of 0.1 was used for each target (130-180 nM) (SI Table 16). At these concentrations, higher sensitivity was observed as compared to larger Fb^0^ values upon displacement by neomycin, a known non-specific RNA-binding small molecule^48^ (SI Table 17). With Tat peptide, RNA concentrations that corresponded to an Fb^0^ value of 0.4 (80-145 nM) were required to obtain acceptable Z’-Factors (SI Table 18). Calculated Z’-Factors were greater than 0.5 for both IDAs, implying that TO-PRO-1 and Tat peptide are reliable and reproducible indicators for these high-throughput screens (SI Tables 19 and 20, SI Figures 5 and 6).^45^

**Figure 4.**
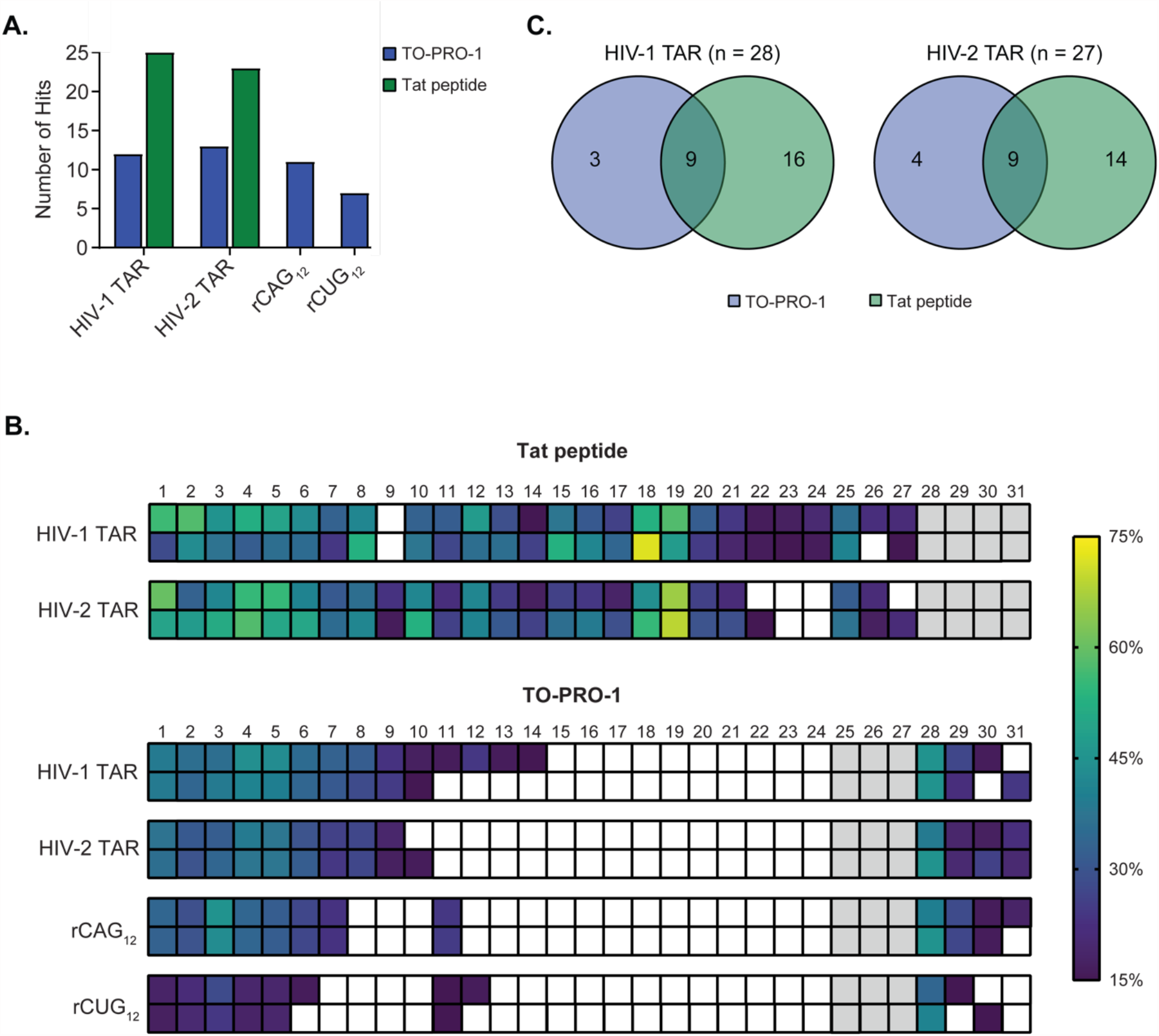
*In vitro* evaluation of the DRTL against RNA targets using TO-PRO-1 and Tat peptide indicator displacement assays. (A) Summary of the number of hits identified for each RNA target. The DRTL was screened against four RNAs using TO-PRO-1 (blue bars) and two RNA targets using Tat peptide (green bars). (B) Heat map representing the percentage displacement of Tat peptide and TO-PRO-1 indicators by small molecule hits. The DRTL was screened at 25 μM in duplicate against the RNAs. A small molecule was considered a hit for an RNA target if 15% or greater displacement of TO-PRO-1 or Tat peptide was observed in both independent measurements. Measurements that were less than 15% displacement are shown in white. Measurements for small molecules that interfered with the emission of an indicator were not considered and are shown in gray. (C) Comparison of the number of hits identified for HIV-1 TAR (left) and HIV-2 TAR (right) RNAs with TO-PRO-1 (blue), Tat peptide (green), or both fluorescent indicators (intersection).

For the TO-PRO-1 IDA, the DRTL was screened at 25 μM against the four RNA targets in duplicate (SI Figures 7-10, SD Table 9). Small molecules were considered “hits” if the signal intensity of TO-PRO-1 was reduced by 15% or greater in duplicate measurements against at least one RNA target. Compounds that enhanced emission of TO-PRO-1 by 15% or greater were excluded from further consideration (n = 76). Multiple hits were identified for each RNA target (Figure 4A), affording hit rates of 0.87% for rCUG12 (n = 7), 1.4% for rCAG_12_ (n = 11), 1.5% for HIV-1 TAR (n = 12), and 1.6% for HIV-2 TAR (n = 13). There was overlap in hits among RNAs as several compounds were found for all four targets (**1**-**5** and **28**), three targets (**6, 7**, and **29**), or two targets (**8, 9, 11**, and **30**), while compounds **10** and **31** were hits for a single RNA (Figure 4B). For most of the hits, the percentage of TO-PRO-1 displaced was within 10% of the displacement for other RNAs, suggesting that these compounds did not have strong binding preferences towards any given target(s) (SI Table 21). Collectively, a total of 15 hits were identified with the TO-PRO-1 IDA. The majority of these hits had scaffolds that mimicked TO-PRO-1 and thus, may bind RNA in a similar manner as the promiscuous indicator (SD Table 10).

For the Tat peptide IDA, the DRTL was also screened at 25 μM against HIV-1 and HIV-2 TAR RNAs in duplicate (SI Figures 11 and 12, SD Table 11). The same criterion was used to define hits (≥ 15% displacement) and small molecules were excluded that interfered with the emission of Tat peptide (n = 97). Multiple hits were identified for each RNA target, with hit rates of 2.9% for HIV-2 TAR (n = 23) and 3.1% for HIV-1 TAR (n = 25) (Figures 4A and 4B). Several of the compounds were hits for both HIV RNA targets (**1-8, 10-21**, and **25**) and had similar Tat peptide displacement percentages for both RNAs (SI Table 22). While some compounds were hits for only HIV-1 TAR (**22-24** and **27**) or HIV-2 TAR (**9** and **26**), the percentage of Tat peptide displacement was usually within 10% of the other RNA target. In total, 27 hits were identified with the Tat peptide IDA. Among these hits, multiple distinct scaffolds that did not bear resemblance to either fluorescent indicator were found (SD Table 10).

Since HIV TAR RNAs were screened in both assays, we directly compared the hits identified from the TO-PRO-1 and Tat peptide IDAs. Collectively, the two IDAs provided a total of 28 hits for HIV-1 TAR and 27 hits for HIV-2 TAR. Only nine of the hits for HIV-1 TAR as well as nine for HIV-2 TAR were found with both indicators (Figure 4C). Differences between the assays were also apparent when considering the number of hits for HIV-1 TAR and HIV-2 TAR RNAs that were identified only with Tat peptide (n = 16 and 14, respectively) as compared to only with TO-PRO-1 (n = 3 and 4, respectively). These differences were partly due to small molecule interference with TO-PRO-1 (**25-27**) or Tat peptide (**28-31**) signal, which excluded a few hits for HIV-1 TAR (n = 4) and HIV-2 TAR (n = 6). Other factors, such as the binding mode, location of small molecule binding relative to each indicator, and the different RNA concentrations used could explain the remaining observed differences. The bias in hits identified by the two screening strategies illustrates the importance of using multiple assays to evaluate RNA:small molecule recognition, especially for primary screens used to prioritize small molecule leads.

### Chemical Space Analysis of DRTL Hits

One of the unique advantages of the DRTL is that it enables a systematic evaluation of R-BIND (SM) chemical space. For example, the RNA target(s) of a hit can be compared to the target(s) reported for the R-BIND (SM) NNs, identifying regions of RNA bioactive space that may be target-specific or target-promiscuous. On the other hand, assessing the distances of hits to their R-BIND (SM) ligand NNs could illuminate boundaries of RNA-privileged chemical space. We also envisioned that DRTL hits could serve as leads for structure-activity relationship studies and biological evaluation in cell-based screens.

The hits identified from our primary screens (n = 31) were NNs to 21 of the 75 R-BIND (SM) ligands (SI Table 23). Approximately half of these R-BIND (SM) ligands target HIV-1 TAR (n = 6) or CUG expanded repeats (n = 7). The remaining R-BIND (SM) NN ligands target a variety of bacterial, mammalian, and viral RNAs. Notably, we found that approximately one third of the hits (**1, 2, 4-7, 9, 27-30**) were NNs to multiple R-BIND (SM) ligands that target a variety of RNAs (SD Table 12). These observations could potentially explain the general lack of *in vitro* binding selectivity of the hits identified in our IDA screens. In the future, we will investigate this hypothesis by screening the DRTL hits to the targets of the R-BIND (SM) NNs and conversely, screening the R-BIND NN ligands to the targets used in this study.

Additionally, we examined the R-BIND-likeness of the hits relative to the entire DRTL. In general, the bin distribution of the hits was similar to the DRTL: the majority of the hits (26/31) were in Bins 6-10 with distances ranging from 1.1961 to 2.1917, and the remaining five compounds were in Bins 2-5 with distances ranging from 0.3981 to 1.0611 (SD Table 12). The similar distribution supports the current distance for library construction, though other variables, such as assay type and ligand screening concentration, contributed to our screening results and consequently, our ability to evaluate the R-BIND-like distance. Several of these variables and future directions are discussed in the Summary and Outlook.

## SUMMARY AND OUTLOOK

A comprehensive understanding and explicit testing of small molecule properties that govern selective RNA-binding and biological activity would significantly expedite the development of chemical probes for RNA. Towards this goal, we curated, analyzed, and evaluated an RNA-focused library that was biased towards 20 cheminformatic properties of bioactive, non-ribosomal RNA-targeting small molecules. The analysis utilized a k-nearest neighbor algorithm^9, 27^ that allowed us to quickly examine > 2 million commercially available small molecules and filter for R-BIND-like ligands. More than 570,000 commercial small molecules were identified to be R-BIND-like and neighbored 61/75 RNA-targeted bioactive ligands. Additionally, the results revealed a lack of commercial compounds in specific regions of R-BIND (SM) chemical space, especially in close proximity to R-BIND (SM) ligands. These limitations illustrate the critical need for theoretical and synthetic development of small molecules that populate unexplored regions of R-BIND (SM) chemical space, and these efforts are currently underway in our laboratory.

This analysis also resulted in the generation of the Duke RNA-Targeted Library (DRTL), one of the largest academic RNA-focused small molecule libraries curated to date (n = 804).^15, 26, 36^ While the cheminformatic and spatial properties of the DRTL represent a subset of R-BIND chemical space, the structural dissimilarity of DRTL to R-BIND ligands led to the identification of several novel RNA-binding small molecules for four therapeutically-relevant RNAs. Small molecule hit rates ranged from 0.87% to 3.1%, which are comparable to reported screens with other RNA-focused libraries.^9, 10, 22^ Comparisons of the hits identified for the same RNA targets revealed differences between the two competition-based assays, with a greater number of hits found using a peptide-based indicator as compared to a small molecule-based indicator under our assay conditions. This direct assay comparison underscores the challenge of screening RNA targets and the need to use multiple assays to identify small molecule leads. Future in-house screens with the DRTL will incorporate and compare methods that directly detect RNA-binding (such as NMR), *in vitro* screens that have a function-related read out, and screens that directly measure biological activity. The DRTL can also be used to easily conduct structure-affinity/activity relationships given the bin-based classification of commercial NN molecules.

We designed the DRTL to gain pivotal insights into RNA:small molecule recognition by systematically evaluating R-BIND (SM) chemical space. The results of our two IDA screens provided data for a preliminary chemical space analysis, which inspired many intriguing hypotheses and future directions. For example, many of the hits from the TO-PRO-1 screen mimicked the chemical architecture of the non-specific, intercalating dye. Additional criterion for library design, such as ligand shape, predicted excitation/emission, or chemical similarity utilizing Tanimoto coefficients, could remove ligands expected to be non-selective prior to purchasing. On the other hand, the lack of selective molecules identified may be due in part to specific assay conditions such as the indicators used and the small molecule concentration. Further, R-BIND (SM) was curated based on biological activity, and few ligands were extensively tested by the scientists for *in vitro* selectivity.^22, 49, 50^ Increasing the number of bioactive RNA ligands, particularly with demonstrated *in vitro* selectivity for structurally similar RNAs and with on-target effects in biological systems, will further refine the boundaries of RNA-privileged chemical space and provide critical benchmarks for RNA-targeted probe design.

These are just a few examples of insights we have already gained from the DRTL to enhance RNA-targeted library design and assay selection. Additional screens of DRTL, utilizing a variety of assays against a diversity of RNA targets, are essential to explicitly evaluate the RNA-privileged small molecule properties and chemical space. We invite members of the community to collaborate with us and use the DRTL in high-throughput, RNA-targeting screens to facilitate these efforts. Further insights into the molecular features that govern selective RNA-binding and biological activity will expedite the discovery of RNA chemical probes — a critical tool for exploring the broad functions and therapeutic potential of RNA.

## Acknowledgements

We thank Dr. Yuqi Zhang and Bilva Sanaba for their contributions of the nearest neighbor algorithm and scripts used to bin as well as randomly select commercial ligands, respectively. Additionally, we thank Kara McCormack and Dr. Terry Hyslop for their assistance with principal component analysis. We thank the Tolbert lab for the preparation of T7 RNA polymerase. We also gratefully thank the Center for HIV-1 RNA Studies and the Duke University Center for RNA Biology for their financial contributions towards the purchase of the DRTL. Many thanks to Dr. So Young Kim and the staff of the Duke University Functional Genomics Facility for their assistance with robotic liquid handling instrumentation that was used to plate and screen the DRTL. Lastly, we thank members of the Hargrove lab for insightful comments and stimulating discussion during the preparation of the manuscript.

## References

1. Morris, K. V.; Mattick, J. S. The rise of regulatory RNA. Nat. Rev. Genet. 2014, 15 (6), 423–437.

2. Connelly, C. M.; Moon, M. H.; Schneekloth, J. S., Jr. The emerging role of RNA as a therapeutic target for small molecules. Cell Chem. Biol. 2016, 23 (9), 1077–1090.

3. Disney, M. D. Targeting RNA with small molecules to capture opportunities at the intersection of chemistry, biology, and medicine. J. Am. Chem. Soc. 2019, 141 (17), 6776–6790.

4. Hermann, T. Small molecules targeting viral RNA. Wiley Interdiscip. Rev. RNA 2016, 7 (6), 726–743.

5. Donlic, A.; Hargrove, A. E. Targeting RNA in mammalian systems with small molecules. Wiley Interdiscip. Rev. RNA 2018, 9 (4), e1477.

6. Moazed, D.; Noller, H. F. Interaction of antibiotics with functional sites in 16S ribosomal RNA. Nature 1987, 327 (6121), 389–394.

7. Lynch, S. R.; Recht, M. I.; Puglisi, J. D. Biochemical and nuclear magnetic resonance studies of aminoglycoside-RNA complexes. Methods Enzymol. 2000, 317,240–61.

8. Hermann, T. Drugs targeting the ribosome. Curr. Opin. Struct. Biol. 2005, 15 (3), 355–366.

9. Morgan, B. S.; Sanaba, B. G.; Donlic, A.; Karloff, D. B.; Forte, J. E.; Zhang, Y.; Hargrove, A. E. R-BIND: An interactive database for exploring and developing RNA-targeted chemical probes. ACS Chem. Biol. 2019, 14 (12), 2691–2700.

10. Donlic, A.; Swanson, E. G.; Chiu, L. Y.; Wicks, S. L.; Juru, A. U.; Cai, Z.; Kassam, K.; Laudeman, C.; Sanaba, B. G.; Sugarman, A.; Han, E.; Tolbert, B. S.; Hargrove, A. E. R-BIND 2.0: An updated database of bioactive RNA-targeting small molecules and associated RNA secondary structures. ACS Chem. Biol. 2022, 17 (6), 1556–1566.

11. Cheung, A. K.; Hurley, B.; Kerrigan, R.; Shu, L.; Chin, D. N.; Shen, Y.; O’Brien, G.; Sung, M. J.; Hou, Y.; Axford, J.; Cody, E.; Sun, R.; Fazal, A.; Fridrich, C.; Sanchez, C. C.; Tomlinson, R. C.; Jain, M.; Deng, L.; Hoffmaster, K.; Song, C.; Van Hoosear, M.; Shin, Y.; Servais, R.; Towler, C.; Hild, M.; Curtis, D.; Dietrich, W. F.; Hamann, L. G.; Briner, K.; Chen, K. S.; Kobayashi, D.; Sivasankaran, R.; Dales, N. A. Discovery of small molecule splicing modulators of Survival Motor Neuron-2 (SMN2) for the treatment of Spinal Muscular Atrophy (SMA). J. Med. Chem. 2018, 61 (24), 11021–11036.

12. Ratni, H.; Ebeling, M.; Baird, J.; Bendels, S.; Bylund, J.; Chen, K. S.; Denk, N.; Feng, Z.; Green, L.; Guerard, M.; Jablonski, P.; Jacobsen, B.; Khwaja, O.; Kletzl, H.; Ko, C. P.; Kustermann, S.; Marquet, A.; Metzger, F.; Mueller, B.; Naryshkin, N. A.; Paushkin, S. V.; Pinard, E.; Poirier, A.; Reutlinger, M.; Weetall, M.; Zeller, A.; Zhao, X.; Mueller, L. Discovery of Risdiplam, a selective Survival of Motor Neuron-2 (SMN2) gene splicing modifier for the treatment of Spinal Muscular Atrophy (SMA). J. Med. Chem. 2018, 61 (15), 6501–6517.

13. Velagapudi, S. P.; Luo, Y.; Tran, T.; Haniff, H. S.; Nakai, Y.; Fallahi, M.; Martinez, G. J.; Childs-Disney, J. L.; Disney, M. D. Defining RNA-small molecule affinity landscapes enables design of a small molecule inhibitor of an oncogenic noncoding RNA. ACS Cent. Sci. 2017, 3 (3), 205–216.

14. Tran, T.; Disney, M. D. Identifying the preferred RNA motifs and chemotypes that interact by probing millions of combinations. Nat. Commun. 2012, 3,1125.

15. Rzuczek, S. G.; Southern, M. R.; Disney, M. D. Studying a drug-like, RNA-focused small molecule library identifies compounds that inhibit RNA toxicity in myotonic dystrophy. ACS Chem. Biol. 2015, 10 (12), 2706–2715.

16. Childs-Disney, J. L.; Tran, T.; Vummidi, B. R.; Velagapudi, S. P.; Haniff, H. S.; Matsumoto, Y.; Crynen, G.; Southern, M. R.; Biswas, A.; Wang, Z. F.; Tellinghuisen, T. L.; Disney, M. D. A massively parallel selection of small molecule-RNA motif binding partners informs design of an antiviral from sequence. Chem. 2018, 4 (10), 2384–2404.

17. Rizvi, N. F.; Santa Maria, J. P., Jr.; Nahvi, A.; Klappenbach, J.; Klein, D. J.; Curran, P. J.; Richards, M. P.; Chamberlin, C.; Saradjian, P.; Burchard, J.; Aguilar, R.; Lee, J. T.; Dandliker, P. J.; Smith, G. F.; Kutchukian, P.; Nickbarg, E. B. Targeting RNA with small molecules: Identification of selective, RNA-binding small molecules occupying drug-like chemical space. SLAS Discov. 2020, 25 (4), 384–396.

18. Haniff, H. S.; Knerr, L.; Liu, X.; Crynen, G.; Bostrom, J.; Abegg, D.; Adibekian, A.; Lekah, E.; Wang, K. W.; Cameron, M. D.; Yildirim, I.; Lemurell, M.; Disney, M. D. Design of a small molecule that stimulates vascular endothelial growth factor A enabled by screening RNA fold-small molecule interactions. Nat. Chem. 2020, 12 (10), 952–961.

19. Yazdani, K.; Jordan, D.; Yang, M.; Fullenkamp, C. R.; Calabrese, D. R.; Boer, R.; Hilimire, T.; Allen, T. E. H.; Khan, R. T.; Schneekloth, J. S., Jr. Machine Learning Informs RNA-Binding Chemical Space. Angew. Chem. Int. Ed. Engl. 2023, 62 (11), e202211358.

20. Santos, R.; Ursu, O.; Gaulton, A.; Bento, A. P.; Donadi, R. S.; Bologa, C. G.; Karlsson, A.; Al-Lazikani, B.; Hersey, A.; Oprea, T. I.; Overington, J. P. A comprehensive map of molecular drug targets. Nat. Rev. Drug Discov. 2017, 16 (1), 19–34.

21. Morgan, B. S.; Forte, J. E.; Culver, R. N.; Zhang, Y.; Hargrove, A. E. Discovery of key physicochemical, structural, and spatial properties of RNA-targeted bioactive ligands. Angew. Chem. Int. Ed. Engl. 2017, 56 (43), 13498–13502.

22. Morgan, B. S.; Forte, J. E.; Hargrove, A. E. Insights into the development of chemical probes for RNA. Nucleic Acids Res. 2018, 46 (16), 8025–8037.

23. Donlic, A.; Morgan, B. S.; Xu, J. L.; Liu, A.; Roble, C., Jr.; Hargrove, A. E. Discovery of small molecule ligands for MALAT1 by tuning an RNA-binding scaffold. Angew. Chem. Int. Ed. Engl. 2018, 57 (40), 13242–13247.

24. Patwardhan, N. N.; Cai, Z.; Umuhire Juru, A.; Hargrove, A. E. Driving factors in amiloride recognition of HIV RNA targets. Org. Biomol. Chem. 2019, 17 (42), 9313–9320.

25. Bodoor, K.; Boyapati, V.; Gopu, V.; Boisdore, M.; Allam, K.; Miller, J.; Treleaven, W. D.; Weldeghiorghis, T.; Aboul-ela, F. Design and implementation of an ribonucleic acid (RNA) directed fragment library. J. Med. Chem. 2009, 52 (12), 3753–3761.

26. Haniff, H. S.; Graves, A.; Disney, M. D. Selective small molecule recognition of RNA base pairs. ACS Comb. Sci. 2018, 20 (8), 482–491.

27. Mitchell, J. B. Machine learning methods in chemoinformatics. Wiley Interdiscip. Rev. Comput. Mol. Sci. 2014, 4 (5), 468–481.

28. Wenderski, T. A.; Stratton, C. F.; Bauer, R. A.; Kopp, F.; Tan, D. S. Principal component analysis as a tool for library design: a case study investigating natural products, brand-name drugs, natural product-like libraries, and drug-like libraries. Methods Mol. Biol. 2015, 1263,225–242.

29. Sauer, W. H.; Schwarz, M. K. Molecular shape diversity of combinatorial libraries: a prerequisite for broad bioactivity. J. Chem. Inf. Comput. Sci. 2003, 43 (3), 987–1003.

30. ChemAxon JChem for Office (Excel), 20.18.0; 2020.

31. Disney, M. D.; Winkelsas, A. M.; Velagapudi, S. P.; Southern, M.; Fallahi, M.; Childs-Disney, J. L. Inforna 2.0: A platform for the sequence-based design of small molecules targeting structured RNAs. ACS Chem. Biol. 2016, 11 (6), 1720–1728.

32. Jakobovits, A.; Smith, D. H.; Jakobovits, E. B.; Capon, D. J. A discrete element 3’ of human immunodeficiency virus 1 (HIV-1) and HIV-2 mRNA initiation sites mediates transcriptional activation by an HIV trans activator. Mol. Cell Biol. 1988, 8 (6), 2555–2561.

33. Todd, P. K.; Paulson, H. L. RNA-mediated neurodegeneration in repeat expansion disorders. Ann. Neurol. 2010, 67 (3), 291–300.

34. Petty, T. J., Bordelon, A. Jason, and Robertson, E. Mary. Thermodynamic characterization of the association of cyanine dyes with DNA. J. Phys. Chem. B 2000, 104 (30), 7221–7227.

35. Asare-Okai, P. N.; Chow, C. S. A modified fluorescent intercalator displacement assay for RNA ligand discovery. Anal. Biochem. 2011, 408 (2), 269–276.

36. Garcia-Lopez, A.; Tessaro, F.; Jonker, H. R. A.; Wacker, A.; Richter, C.; Comte, A.; Berntenis, N.; Schmucki, R.; Hatje, K.; Petermann, O.; Chiriano, G.; Perozzo, R.; Sciarra, D.; Konieczny, P.; Faustino, I.; Fournet, G.; Orozco, M.; Artero, R.; Metzger, F.; Ebeling, M.; Goekjian, P.; Joseph, B.; Schwalbe, H.; Scapozza, L. Targeting RNA structure in SMN2 reverses spinal muscular atrophy molecular phenotypes. Nat. Commun. 2018, 9 (1), 2032.

37. Ursu, A.; Wang, K. W.; Bush, J. A.; Choudhary, S.; Chen, J. L.; Baisden, J. T.; Zhang, Y. J.; Gendron, T. F.; Petrucelli, L.; Yildirim, I.; Disney, M. D. Structural features of small molecules targeting the RNA repeat expansion that causes genetically defined ALS/FTD. ACS Chem. Biol. 2020, 15 (12), 3112–3123.

38. Wicks, S. L.; Hargrove, A. E. Fluorescent indicator displacement assays to identify and characterize small molecule interactions with RNA. Methods 2019, 167,3–14.

39. Weeks, K. M.; Crothers, D. M. RNA recognition by Tat-derived peptides: Interaction in the major groove? Cell 1991, 66 (3), 577–588.

40. Calnan, B. J.; Tidor, B.; Biancalana, S.; Hudson, D.; Frankel, A. D. Arginine-mediated RNA recognition: The arginine fork. Science 1991, 252 (5009), 1167–1171.

41. Matsumoto, C.; Hamasaki, K.; Mihara, H.; Ueno, A. A high-throughput screening utilizing intramolecular fluorescence resonance energy transfer for the discovery of the molecules that bind HIV-1 TAR RNA specifically. Bioorg. Med. Chem. Lett. 2000, 10 (16), 1857–1861.

42. Patwardhan, N. N.; Cai, Z.; Newson, C. N.; Hargrove, A. E. Fluorescent peptide displacement as a general assay for screening small molecule libraries against RNA. Org. Biomol. Chem. 2019, 17 (7), 1778–1786.

43. Abulwerdi, F. A.; Shortridge, M. D.; Sztuba-Solinska, J.; Wilson, R.; Le Grice, S. F.; Varani, G.; Schneekloth, J. S., Jr. Development of small molecules with a noncanonical binding mode to HIV-1 Trans Activation Response (TAR) RNA. J. Med. Chem. 2016, 59 (24), 11148–11160.

44. Leamy, K. A.; Yennawar, N. H.; Bevilacqua, P. C. Cooperative RNA folding under cellular conditions arises from both tertiary structure stabilization and secondary structure destabilization. Biochemistry 2017, 56 (27), 3422–3433.

45. Zhang, J. H.; Chung, T. D.; Oldenburg, K. R. A simple statistical parameter for use in evaluation and validation of high throughput screening assays. J. Biomol. Screen. 1999, 4 (2), 67–73.

46. Boger, D. L.; Fink, B. E.; Brunette, S. R.; Tse, W. C.; Hedrick, M. P. A simple, highresolution method for establishing DNA binding affinity and sequence selectivity. J. Am. Chem. Soc. 2001, 123 (25), 5878–5891.

47. Del Villar-Guerra, R.; Gray, R. D.; Trent, J. O.; Chaires, J. B. A rapid fluorescent indicator displacement assay and principal component/cluster data analysis for determination of ligand-nucleic acid structural selectivity. Nucleic Acids Res. 2018, 46 (7), e41.

48. Thomas, J. R.; Hergenrother, P. J. Targeting RNA with small molecules. Chem. Rev. 2008, 108 (4), 1171–1224.

49. Kelly, M. L.; Chu, C. C.; Shi, H.; Ganser, L. R.; Bogerd, H. P.; Huynh, K.; Hou, Y.; Cullen, B. R.; Al-Hashimi, H. M. Understanding the characteristics of nonspecific binding of druglike compounds to canonical stem-loop RNAs and their implications for functional cellular assays. RNA 2021, 27 (1), 12–26.

50. Bush, J. A.; Williams, C. C.; Meyer, S. M.; Tong, Y.; Haniff, H. S.; Childs-Disney, J. L.; Disney, M. D. Systematically studying the effect of small molecules interacting with RNA in cellular and preclinical models. ACS Chem. Biol. 2021, 16 (7), 1111–1127.

