## Supplementary Information for "Probing Bioactive Chemical Space to Discover RNA-Targeted Small Molecules"

### SI-1. Cheminformatic Calculations

SMILES strings for all molecules were batch-processed. Using the ChemAxon Calculator Plugins, ligands were corrected to their major protonation and tautomeric states (pH = 7.4), and then the 20 cheminformatic parameters were calculated using the ChemAxon Chemical Terms Evaluator (Marvin 16.4.25.0, 2016, http://www.chemaxon.com) (SI Table 1).[1]

**SI Table 1**: Cheminformatic descriptors

| Category | Parameter | Description | Chemical Terms Evaluator Expression |
|---|---|---|---|
| Medicinal chemistry | MW | Molecular weight | mass() |
| | HBA | Number of hydrogen bond acceptors | acceptorCount() |
| | HBD | Number of hydrogen bond donors | donorCount() |
| | LogP | n-Octanol/Water partition coefficient | logPKLOP() |
| | RotB | Number of rotatable bonds | rotatableBondCount() |
| | tPSA | Topological polar surface area | PSA() |
| | LogD | n-Octanol/Water distribution coefficient | logD('7.4') |
| Structural | N | Number of nitrogen atoms | atomCount('7') |
| | O | Number of oxygen atoms | atomCount('8') |
| | Rings | Number of rings | ringCount() |
| | ArRings | Number of aromatic rings | aromaticRingCount() |
| | HetRings | Number of heteroatom-containing rings | heteroRingCount() |
| | SysRings | Number of ring systems | ringSystemCount() |
| | SysRR | Ring complexity | ringCount()/ringSystemCount() |
| Molecular complexity & recognition | $Fsp^3$ | Fraction of $sp^3$ hybridized carbons | Count(filter('atno()==6&&connections()==4'))/atomCount('6') |
| | nStereo | Number of stereocenters | chiralCenterCount() |
| | ASA | Accessible surface area | ASA() |
| | relPSA | Relative polar surface area | PSA()/vanDerWaalsSurfaceArea() |
| | TC | Total charge | totalCharge() |
| | VWSA | van der Waals surface area | vanDerWaalsSurfaceArea() |

### SI-2. Nearest Neighbor Search

A k-Nearest neighbors algorithm was used, where k = 1, to evaluate overlap between pairs of small molecules.[2, 3] The nearest neighbor distance for R-BIND (SM) v1.1 was calculated by: (1) Normalizing each cheminformatic parameter of every small molecule to the mean and standard deviation of the descriptor for the library; (2) Using the normalized values to calculate multi-dimensional distances in space between each pair of R-BIND (SM) ligands; (3) Averaging the smallest nearest neighbor distance (d) for each R-BIND (SM); and (4) Mapping each R-BIND (SM) as a multi-dimensional sphere with a radius of d. Similarly, commercial ligands from ChemBridge (n = 1,146,666) and ChemDiv (n = 1,465,277) were normalized to the R-BIND (SM) v1.1 library and mapped as a multi-dimensional sphere with a radius of the smallest averaged nearest neighbor distance. A commercial ligand was considered "R-BIND-like" if its multi-dimensional sphere overlapped with any R-BIND (SM) multi-dimensional sphere.

**SI Table 2:** Distribution of commercial small molecule (SM) nearest neighbors (NN) for R-BIND (SM) ligands (n = 61)

| Measure | # of Commercial SM NN |
|---:|---:|
| Minimum | 1 |
| First Quartile | 22 |
| Median | 1057 |
| Average | 17968 |
| Third Quartile | 15432 |
| Maximum | 197433 |

**SI Table 3:** R-BIND (SM) ligands that had no commercial small molecule nearest neighbors

| R-BIND (SM) v1.1 | Shortest Distance to a Commercial SM |
|---:|---:|
| 0004 | 3.6506 |
| 0014 | 4.4924 |
| 0020 | 2.2717 |
| 0022 | 2.8763 |
| 0023 | 3.5601 |
| 0024 | 2.4859 |
| 0044 | 3.2542 |
| 0045 | 3.1083 |
| 0046 | 3.0247 |
| 0055 | 2.3231 |
| 0069 | 3.4768 |
| 0071 | 3.7316 |
| 0072 | 3.0493 |
| 0073 | 3.5122 |

**SI Table 4:** Number of R-BIND (SM) nearest neighbors (NN) to those that had no commercial ligands within the shortest average distance (d)

| R-BIND (SM) v1.1 | Shortest Distance to an R-BIND (SM) | # of R-BIND (SM) NN (d ≤ 2.1932) |
|---:|---:|---:|
| 0004 | 6.1207 | - |
| 0014 | 7.3373 | - |
| 0020 | 2.1858 | 1 |
| 0022 | 2.5842 | - |
| 0023 | 1.8107 | 1 |
| 0024 | 4.6910 | - |
| 0044 | 0.3453 | 2 |
| 0045 | 0.3453 | 2 |
| 0046 | 2.8889 | - |
| 0055 | 2.5842 | - |
| 0069 | 2.2153 | - |
| 0071 | 3.6357 | - |
| 0072 | 1.8107 | 1 |
| 0073 | 3.6707 | - |

### SI-3. Curation of the Duke RNA-Targeted Library

All “R-BIND-like” commercial ligands for a given R-BIND (SM) were sorted according to the nearest neighbor distance between the pair into one of ten bins. The range of nearest neighbor distances for each bin was 0-0.21923 (Bin 1), 0.21924-0.43846 (Bin 2), 0.43847-0.65769 (Bin 3), 0.65770-0.87692 (Bin 4), 0.87693-1.0962 (Bin 5), 1.0963-1.3154 (Bin 6), 1.3155-1.5346 (Bin 7), 1.5347-1.7538 (Bin 8), 1.7539-1.9731 (Bin 9), and 1.9732-2.1923 (Bin 10). For every R-BIND (SM), three commercial ligands were randomly selected from each bin, if possible, and no ligand could be selected twice. In total, 435 and 369 compounds were purchased from ChemBridge and ChemDiv, respectively. Each compound (up to 9.5 μmol) was manually dissolved to 10 mM in DMSO and aliquots were robotically transferred to 384-well plates. Parent and daughter plates were stored at -80 ºC.

### SI-4. Cheminformatic Comparisons Between Libraries

**Mann-Whitney U Test**
All cheminformatic statistical comparisons between libraries were performed using an independent two-group Mann-Whitney U test in R statistical software (v3.4.3, 2016).

**SI Table 5:** Statistical comparison of DRTL and R-BIND (SM) v1.1 descriptors

| Parameter | Means | | *P* Value |
|---|---|---|---|
| | DRTL | R-BIND (SM) v1.1 | |
| MW | 347 | 366 | 0.156 |
| HBA | 3.87 | 3.89 | 0.849 |
| HBD | 1.61 | 2.55 | < 0.001 |
| LogP | 2.23 | 1.65 | 0.010 |
| RotB | 4.18 | 4.79 | 0.500 |
| tPSA | 75.3 | 82.2 | 0.104 |
| LogD | 1.87 | 0.31 | < 0.001 |
| N | 4.09 | 4.49 | 0.026 |
| O | 1.86 | 1.68 | 0.097 |
| Rings | 3.48 | 3.73 | 0.128 |
| ArRings | 2.84 | 3.04 | 0.246 |
| HetRings | 2.09 | 2.12 | 0.894 |
| SysRings | 2.59 | 2.47 | 0.254 |
| SysRR | 1.48 | 1.81 | 0.021 |
| $Fsp^3$ | 0.25 | 0.28 | 0.289 |
| nStereo | 0.13 | 0.32 | 0.028 |
| ASA | 565 | 601 | 0.052 |
| relPSA | 0.17 | 0.18 | 0.935 |
| TC | 0.31 | 1.01 | < 0.001 |
| VWSA | 485 | 536 | 0.006 |

(light shading) $P < 0.05$; (dark shading) $P < 0.001$

**SI Table 6:** Statistical comparison of DRTL and R-BIND (SM) neighbors (n = 61) descriptors

| | Means | | |
|---|---|---|---|
| **Parameter** | **DRTL** | **R-BIND (SM) v1.1** | ***P* Value** |
| MW | 347 | 352 | 0.869 |
| HBA | 3.87 | 3.97 | 0.666 |
| HBD | 1.61 | 2.31 | < 0.001 |
| LogP | 2.23 | 1.74 | 0.029 |
| RotB | 4.18 | 4.46 | 0.825 |
| tPSA | 75.3 | 78.9 | 0.411 |
| LogD | 1.87 | 0.79 | < 0.001 |
| N | 4.09 | 4.34 | 0.206 |
| O | 1.86 | 1.62 | 0.076 |
| Rings | 3.48 | 3.61 | 0.405 |
| ArRings | 2.84 | 2.93 | 0.662 |
| HetRings | 2.09 | 2.08 | 0.972 |
| SysRings | 2.59 | 2.46 | 0.226 |
| SysRR | 1.48 | 1.70 | 0.030 |
| $Fsp^3$ | 0.25 | 0.27 | 0.521 |
| nStereo | 0.13 | 0.16 | 0.357 |
| ASA | 565 | 585 | 0.264 |
| relPSA | 0.17 | 0.18 | 0.966 |
| TC | 0.31 | 0.79 | < 0.001 |
| VWSA | 485 | 512 | 0.190 |

(light shading) $P < 0.05$; (dark shading) $P < 0.001$

**Principal Component Analysis**

Principal component analysis (PCA) on centered and scaled cheminformatic parameters of R-BIND (SM) ligands was performed using the prcomp function in the RStudio open-source statistical computing software package (v1.4.1106).[1] Small molecules in the DRTL were treated as supplemental observations and did not contribute to the principal components. Using the predict function, coordinates for the DRTL were calculated by: (1) Normalizing each cheminformatic parameter for every supplemental observation to the mean and standard deviation of the descriptor for the R-BIND (SM) library and (2) Multiplying the normalized values with the eigenvectors of the principal components.

**Principal Moments of Inertia Calculations**

Using protonation- and tautomer-corrected SMILES strings, low energy conformations of each molecule were calculated using the Conformation Search algorithm in the Molecular Operating Environment (MOE, v2019.01) software package. The Conformation Search function was performed using the stochastic method with the MMFF94 force field and generalized Born solvation model. The input for each parameter is listed in SI Table 7, and the following options were checked: 1) calculate force field partial charges and 2) hydrogens. Following conformational search, the normalized principal moments of inertia descriptors, *npr1* and *npr2*, were computed for each conformer in MOE. The Boltzmann weighted average for *npr1* and *npr2* of each molecule was calculated by using Equation (1):

$$A_i = \frac{\sum_i A e^{-\frac{E_i}{k_B T}}}{\sum_i e^{-\frac{E_i}{k_B T}}} \quad (1)$$

where $A$ is the calculated *npr1* or *npr2* value of the conformation, $E_i$ is the relative energy of the conformation (kcal/mol), $k_B$ is the Boltzmann constant (0.001986 kcal/(mol*K)), and T is the temperature (298 K). The resulting coordinates were plotted on a triangular graph where the vertices represent rod- (0,1), sphere- (1,1), or disc-like (0.5, 0.5) shape.

**SI Table 7:** Parameters for conformation search

| Parameter | Input |
|---|---|
| Rejection limit | 100 |
| Iteration limit | 10000 |
| RMS gradient | 0.005 |
| MM iteration limit | 500 |
| RMSD limit | 0.15 |
| Energy window | 3 |
| Conformation limit | 10000 |

**SI Table 8:** Average normalized principal moment of inertia for each library

| Library | $I_1/I_3$ | $I_2/I_3$ |
|---|---|---|
| R-BIND (SM) v1.1 | 0.22 | 0.85 |
| DRTL | 0.25 | 0.85 |

**Cell-based Partitioning**

In Python (Python Language Reference v2.7, http://www.python.org), cell-based partitioning was performed. The principal moments of inertia triangle was defined by three lines: y = 1; y = -x + 1; and y = x. Then, the triangular graph was partitioned into four isosceles triangles of equal size by defining the slopes and area of each sub-triangle. The Boltzmann weighted averages for *npr1* and *npr2* of each molecule were rounded to three significant figures, and if the coordinate fell within a partition, the identity of the triangle was returned.

**SI Table 9:** Small molecule counts for each library in the four triangle partitions

| Library | Rod | Hybrid | Sphere | Disc | Total |
|---|---|---|---|---|---|
| R-BIND (SM) v1.1 | 56 | 6 | 0 | 13 | 75 |
| DRTL | 574 | 113 | 1 | 116 | 804 |

**Kolmogorov-Smirnov Test**

Rod, disc, and sphere statistical comparisons between libraries were performed using a two-sided, non-parametric Kolmogorov-Smirnov test in R statistical software (v4.0.4, 2021).

**SI Table 10:** Statistical comparisons of the distributions from the rod, disc, and sphere vertices for DRTL and R-BIND (SM)

| | ***P* values** |
|---|---|
| **Shape** | **DRTL / R-BIND (SM) v1.1** |
| Rod | 0.006 |
| Disc | 0.001 |
| Sphere | 0.026 |

**SI Table 11:** Statistical comparisons of the distributions from the rod, disc, and sphere vertices for DRTL and R-BIND (SM) neighbors (n = 61)

| | ***P* values** |
|---|---|
| **Shape** | **DRTL / R-BIND (SM) Neighbors** |
| Rod | 0.023 |
| Disc | 0.081 |
| Sphere | 0.005 |

**Cumulative Distribution**

The Euclidean distance of each small molecule from the rod (0,1), sphere (1,1), and disc (0.5, 0.5) vertices was calculated. The distances to a vertex for each library were ordered from shortest to longest, and the cumulative frequency for each distance was calculated. Cumulative distance distribution graphs were generated using GraphPad Prism (version 9.0.1 for Windows, GraphPad Software, La Jolla California USA, www.graphpad.com).

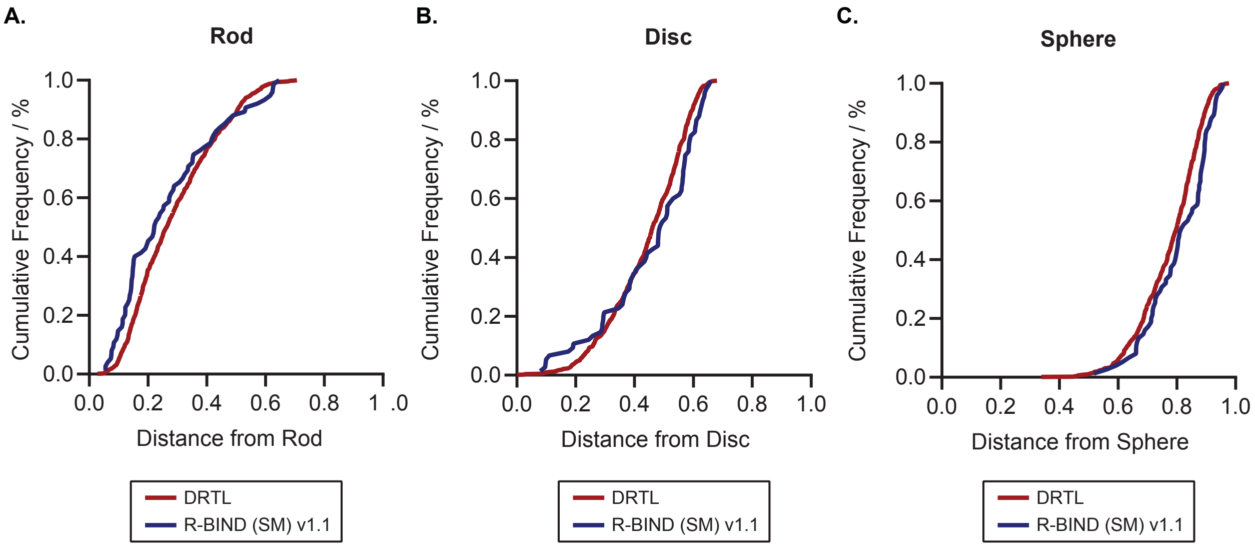

**SI Figure 1:** Cumulative distribution of the distance from the a) rod, b) disc, and c) sphere vertices for DRTL and R-BIND (SM) v1.1.

**Tanimoto Dissimilarity Scores**

The JCDissimilarityCFTanimoto function in JChem for Office (Excel), version 20.18.0 (ChemAxon; http://www.chemaxon.com) was used to calculate the Tanimoto dissimilarity score between small molecules.

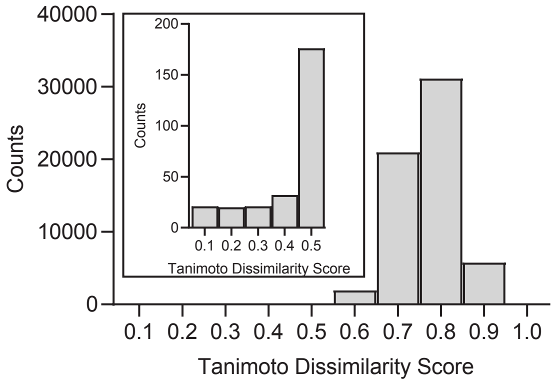

**SI Figure 2:** Tanimoto dissimilarity scores for pairwise comparisons between the DRTL (n = 804) and R-BIND (SM) v1.1 (n = 75). A score of 1.0 indicates that two ligands are highly dissimilar. The inset shows a zoom in of the counts for Tanimoto dissimilarity scores of 0.1-0.5.

### SI-5. General Experimental Information

All buffers were prepared with molecular biology grade reagents and nuclease-free water, then filtered using a 0.2 μM polyethersulfone filter prior to use. To avoid RNase contamination, surfaces were cleaned with RNaseZap prior to RNA handling. TO-PRO-1 was purchased from Thermo Fisher Scientific (T3602). Fluorescently labeled Tat peptide [N-(5-FAM)-AAARKKRRQRRRAAAK-(TAMRA)-C] was purchased from LifeTein and used without further purification. Neomycin was purchased from Amresco and used without further purification. Low volume 384-well plates were purchased from Corning (4514). A BioTek Multiflo microplate dispenser, that was pre-treated with 20 mg/mL of poly(vinylpolypyrrolidone) for 10 minutes to prevent adsorption of assay reagents to the dispenser tubes, was used to robotically plate aliquots of RNA and fluorescent indicator for small molecule screening. Aliquots of small molecules were robotically transferred using an Echo 550 Acoustic liquid handler (Labcyte). All fluorescence measurements were recorded using a CLARIOstar Plus plate reader (BMG Labtech).

### SI-6. RNA Preparation

All RNAs were synthesized through *in vitro* transcription. Sense DNA templates carrying the T7 promoter and complementary antisense strands containing two 2'-O-methylated residues at the 5'-end (SI Table 12) were purchased as solids from Integrated DNA Technologies (IDT). Sense and antisense DNA strands were annealed. Specifically, 25 µM of sense and antisense DNA templates were mixed with 3 mM $MgCl_2$, the mixture was heated at 95 ºC for 10 minutes and snap-cooled on ice for at least one hour. Annealed DNA templates were confirmed by running an aliquot of annealed DNA on a Novex® DNA retardation gel. Each annealed double-stranded DNA template (0.4 µM) was mixed with 90.9 mM Tris (pH 7.4), 22.7 mM $MgCl_2$, 27.3 mM dithiothreitol, 1.8 mM spermidine, 2.7 mM rNTP (rATP, rCTP, rGTP, and rUTP), and 2.7 U/µL T7 RNA polymerase (custom preparation). Mixtures were heated at 37 ºC for at least 12 hours. RNAs (SI Table 13) were purified by denaturing gel electrophoresis (20% polyacrylamide [29:1 acrylamide/bis-acrylamide], 8 M urea). RNAs were eluted in 50 mM Tris, 300 mM NaOAc, 10 mM EDTA, pH 7.2. RNAs were concentrated using a 3 KDa molecular weight cutoff Amicon® filtration membrane, washed extensively with nuclease-free water, then isolated by ethanol precipitation. RNA purity was confirmed by running an aliquot of isolated RNA on a 15% Novex® TBE-Urea gel. Concentrations of RNA were determined using Beer's law with the following extinction coefficients of 268,900 $M^{-1}$ $cm^{-1}$ (HIV-1 TAR), 269,200 $M^{-1}$ $cm^{-1}$ (HIV-2 TAR), 386,100 $M^{-1}$ $cm^{-1}$ ($rCAG_{12}$), and 334,500 $M^{-1}$ $cm^{-1}$ ($rCUG_{12}$). All RNAs were annealed prior to use. For HIV-1 and HIV-2 TAR RNAs, RNAs were diluted to 25 µM in nuclease-free water, heated at 95 ºC for 10 minutes, and snap-cooled on ice for at least one hour. For $rCAG_{12}$ and $rCUG_{12}$ RNAs, RNAs were diluted to 3 µM in nuclease-free water, heated at 95 ºC for 10 minutes, and cooled on bench top for at least one hour.

**SI Table 12**: Sequences of DNA constructs

| DNA | Sequences |
|---|---|
| HIV-1 TAR | 5'-GCAGCTAATACGACTCACTATAGGCAGATCTGAGCCTGGGAGCTCTCTGCC-3'<br>3'-CGTCGATTATGCTGAGTGATATCCGTCTAGACTCGGACCCTCGAGAGACmG*mG*-5' |
| HIV-2 TAR | 5'-GCAGCTAATACGACTCACTATAGGAGATTGAGCCCTGGGAGGTTCTCTCC-3'<br>3'-CGTCGATTATGCTGAGTGATATCCTCTAACTCGGGACCCTCCAAGAGAmG*mG*-5' |
| $rCAG_{12}$ | 5'-GCAGCTAATACGACTCACTATAGGCAGCAGCAGCAGCAGCAGCAGCAGCAGCAGCAGCAGCC-3'<br>3'-CGTCGATTATGCTGAGTGATATCCGTCGTCGTCGTCGTCGTCGTCGTCGTCGTCGTCGTCmG*mG*-5' |
| $rCUG_{12}$ | 5'-GCAGCTAATACGACTCACTATAGGCTGCTGCTGCTGCTGCTGCTGCTGCTGCTGCTGCTGCC-3'<br>3'-CGTCGATTATGCTGAGTGATATCCGACGACGACGACGACGACGACGACGACGACGACGACmG*mG*-5' |

* indicates 2'-OMe modified nucleotide

**SI Table 13**: Sequences of RNA constructs

| RNA | Sequence (5' to 3') |
|---|---|
| HIV-1 TAR | GGCAGAUCUGAGCCUGGGAGCUCUCUGCC |
| HIV-2 TAR | GGAGAUUGAGCCCUGGGAGGUUCUCUCC |
| $rCAG_{12}$ | GGCAGCAGCAGCAGCAGCAGCAGCAGCAGCAGCAGCAGCC |
| $rCUG_{12}$ | GGCUGCUGCUGCUGCUGCUGCUGCUGCUGCUGCUGCUGCC |

### SI-7. Optimization of Indicator Displacement Assays

**RNA Titrations with TO-PRO-1**

In a 384-well plate, samples (20 µL) were manually plated, containing 500 nM TO-PRO-1, 140 mM KCl, 10 mM $NaH_2PO_4$, 1 mM $MgCl_2$, 0.1 mM EDTA, 0.01% Triton-x-100, and 5% DMSO (pH 7.2). In a 96-well plate, RNAs were serially diluted in buffer by 3-fold and transferred to the 384-well plate, resulting in one of the following concentrations of RNA: 0.0219, 0.0343, 0.0658, 0.102, 0.200, 0.310, 0.590, 0.930, 1.78, 2.78, 5.33, 8.33, 16.0, or 25.0 µM. The plate was placed on a shaker at 100 rpm for 5 mins, centrifuged at 4000 rpm for 1 min, and incubated at ambient temperature in the dark for 30 mins. The plate was excited at 492 nm and emission was recorded at 575 nm with 9 and 15 nm slits, respectively. A read height of 11.3 mm was used for all measurements. The gain was adjusted for each RNA to 75% of maximum fluorescence of samples containing 25 µM RNA: 1708 (HIV-1 TAR), 1654 (HIV-2 TAR), 1881 ($rCAG_{12}$), and 1964 ($rCUG_{12}$). Binding isotherms were generated by plotting the fluorescence emission intensity vs. [RNA]. Apparent dissociation constants ($K_{d(app)}$) were calculated in GraphPad Prism (version 7.04 for Windows, GraphPad Software, San Diego, California USA, www.graphpad.com) by fitting to the One Site Binding (hyperbola) model. Titrations were conducted in three inter-run replicates and each replicate contained three technical replicates.

**SI Table 14:** Apparent dissociation constant of TO-PRO-1 for RNA targets

| Measure | HIV-1 TAR | HIV-2 TAR | $rCAG_{12}$ | $rCUG_{12}$ |
|---|---|---|---|---|
| $K_{d(app)}$ (µM)[a]: | 0.94 ± 0.07 | 0.83 ± 0.04 | 0.71 ± 0.04 | 1.18 ± 0.07 |

[a]Values represent the fit of three inter-run replicates, where each inter-run replicate consisted of three technical replicates.

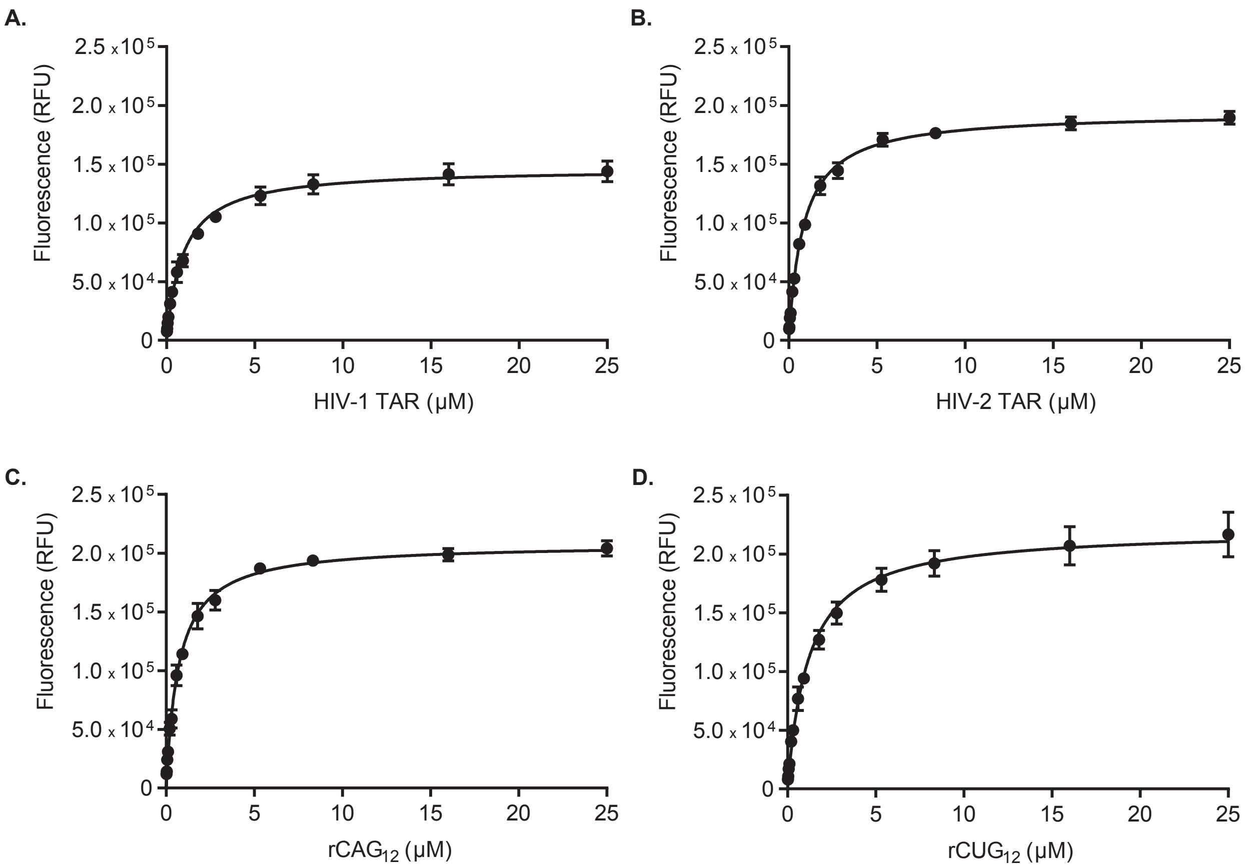

**SI Figure 3.** Titration of HIV-1 TAR (A), HIV-2 TAR (B), rCAG$_{12}$ (C), and rCUG$_{12}$ (D) (0.022-25 μM) at a constant concentration of TO-PRO-1 (500 nM) in 140 mM KCl, 10 mM $NaH_2PO_4$, 1 mM $MgCl_2$, 0.1 mM EDTA, 0.01% Triton-x-100, 5% DMSO, pH 7.2. Samples were excited at 492 nm and emission was recorded at 575 nm. Error bars represent standard deviation between three inter-run replicates. Each inter-run replicate consisted of three technical replicates. Apparent dissociation constants ($K_{d(app)}$) were calculated in GraphPad Prism by fitting to the One Site Binding (hyperbola) model.

**RNA Titrations with Tat Peptide**

In a 384-well plate, samples (16 μL) were manually plated, containing 50 nM Tat peptide, 140 mM KCl, 10 mM $NaH_2PO_4$, 1 mM $MgCl_2$, 0.1 mM EDTA, 0.01% Triton-x-100, and 5% DMSO (pH 7.2). In a 96-well plate, RNAs were serially diluted in buffer by 1.75-fold and transferred to the 384-well plate. Final RNA concentrations were: 0.00, 0.0071, 0.0125, 0.0218, 0.0382, 0.0668, 0.117, 0.205, 0.358, 0.627, 1.10, 1.92, 3.36, 5.88, 10.3, and 18.0 μM for HIV-1 TAR and 0.00, 0.0040, 0.0069, 0.0121, 0.0212, 0.0371, 0.0650, 0.114, 0.199, 0.348, 0.609, 1.07, 1.87, 3.27, 5.71, and 10.0 μM for HIV-2 TAR. The plate was placed on a shaker at 100 rpm for 2 mins, centrifuged at 4000 rpm for 1 min, and incubated at ambient temperature in the dark for 30 mins. The plate was excited at 485 nm and emission was recorded at 590 nm with 15 nm slits. A read height of 11.5 mm was used for all measurements. The gain was adjusted for each RNA to 80% of maximum fluorescence of samples containing the highest RNA concentration: 2237 (HIV-1 TAR) and 2305 (HIV-2 TAR). Binding isotherms were generated by plotting the fluorescence emission intensity vs. [RNA]. Apparent dissociation constants ($K_{d(app)}$) were calculated in GraphPad Prism (version 7.04 for Windows, GraphPad Software, San Diego, California USA, www.graphpad.com) by fitting to the One site – Total binding model. Titrations were conducted in three inter-run replicates and each replicate contained three technical replicates.

**SI Table 15:** Apparent dissociation constant of Tat peptide for RNA targets

| Measure | HIV-1 TAR | HIV-2 TAR |
|---|---|---|
| $K_{d(app)}$ (nM)[a]: | 188 ± 22 | 92 ± 6 |

[a]Values represent the fit of three inter-run replicates, where each inter-run replicate consisted of three technical replicates.

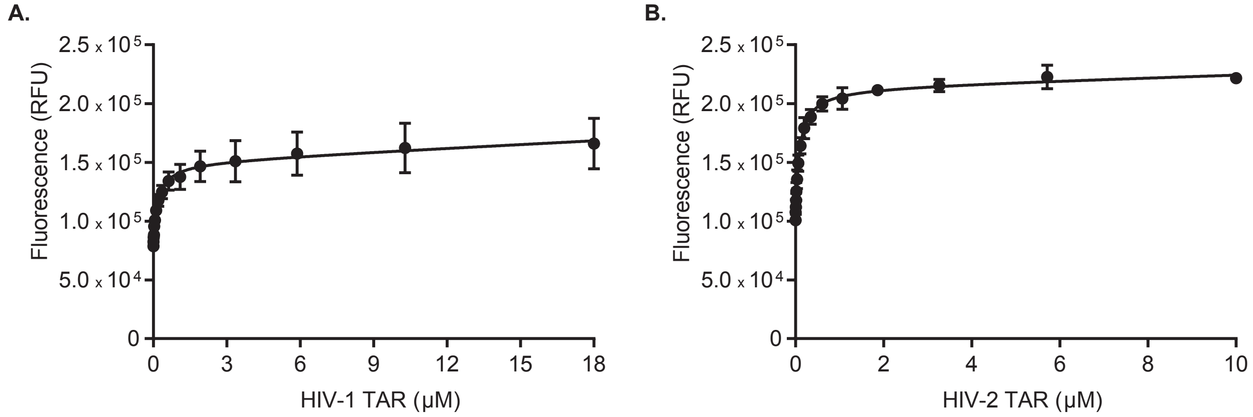

**SI Figure 4.** Titration of RNA at a constant concentration of Tat peptide (50 nM) in 140 mM KCl, 10 mM $NaH_2PO_4$, 1 mM $MgCl_2$, 0.1 mM EDTA, 0.01% Triton-x-100, 5% DMSO, pH 7.2. RNA concentrations for HIV-1 (A) and HIV-2 TAR (B) were 0-18 µM and 0-10 µM, respectively. Samples were excited at 485 nm and emission was recorded at 590 nm. Error bars represent standard deviation between three inter-run replicates. Each inter-run replicate consisted of three technical replicates. In one technical replicate for HIV-2 TAR, fluorescence measurements at 5.7 and 10 µM were found to be significant outliers by Grubbs test and were removed. Apparent dissociation constants ($K_{d(app)}$) were calculated in GraphPad Prism by fitting to the One site – Total binding model.

**Initial Fraction of Bound Indicator Calculations**

The initial fraction bound ($Fb^0$) of TO-PRO-1 and Tat peptide as a function of the binding constant of each RNA was calculated using reported methods.[4]

**SI Table 16:** Initial fraction ($Fb^0$) of TO-PRO-1 bound at respective RNA concentrations

| RNA | $K_{d(app)}$ (μM)[a] | [RNA] (μM) | Initial saturation ($Fb^0$)[b] |
|---|---|---|---|
| HIV-1 TAR | 0.94 ± 0.07 | 0.15 | 0.10 |
| HIV-2 TAR | 0.83 ± 0.04 | 0.14 | 0.10 |
| $rCAG_{12}$ | 0.71 ± 0.04 | 0.13 | 0.10 |
| $rCUG_{12}$ | 1.18 ± 0.07 | 0.18 | 0.10 |

[a]Values represent the fit of three inter-run replicates, where each inter-run replicate consisted of three technical replicates. [b]Initial saturation fraction was calculated using the corresponding $K_{d(app)}$ value, RNA concentration, and 0.5 μM TO-PRO-1

**SI Table 17:** Percentage displacement of TO-PRO-1 (0.5 μM) fluorescence by Neomycin (5 μM) at different concentrations of RNA

| RNA | [RNA] (μM) | Initial saturation ($Fb^0$) | Displacement (%)[a] |
|---|---|---|---|
| HIV-1 TAR | 0.15 | 0.10 | 70 |
| | 0.33 | 0.20 | 65 |
| | 0.55 | 0.30 | 64 |
| $rCUG_{12}$ | 0.18 | 0.10 | 60 |
| | 0.97 | 0.40 | 46 |

[a]Values represent the average of three technical replicates

**SI Table 18:** Initial fraction ($Fb^0$) of Tat peptide bound at respective RNA concentrations

| RNA | $K_{d(app)}$ (nM)[a] | [RNA] (nM) | Initial saturation ($Fb^0$)[b] |
|---|---|---|---|
| HIV-1 TAR | 188 ± 22 | 145 | 0.40 |
| HIV-2 TAR | 92 ± 6 | 80 | 0.40 |

[a]Values represent the fit of three inter-run replicates, where each inter-run replicate consisted of three technical replicates. [b]Initial saturation fraction was calculated using the corresponding $K_{d(app)}$ value, RNA concentration, and 50 nM Tat peptide

**Z'-Factor Measurements for TO-PRO-1 Indicator Displacement Assay**

In a 384-well plate, 96 positive controls (20 µL) were manually plated in alternating columns, containing 500 nM TO-PRO-1, 140 mM KCl, 10 mM $NaH_2PO_4$, 1 mM $MgCl_2$, 0.1 mM EDTA, 0.01% Triton-x-100, and 5% DMSO (pH 7.2). Similarly, 96 negative controls (20 µL) were manually plated in alternating columns, containing the same reagents and additionally, RNA at a concentration that corresponded to an $Fb^0$ of 0.1; 0.15 µM (HIV-1 TAR), 0.14 µM (HIV-2 TAR), 0.13 µM ($rCAG_{12}$), or 0.18 µM ($rCUG_{12}$). The plate was placed on a shaker at 100 rpm for 5 mins, centrifuged at 4000 rpm for 1 min, and incubated at ambient temperature in the dark for 30 mins. The plate was excited at 492 nm and emission was recorded at 575 nm with 9 and 15 nm slits, respectively. A read height of 11.3 mm and gain of 1708 (HIV-1 TAR), 1654 (HIV-2 TAR), 1881 ($rCAG_{12}$), or 1964 ($rCUG_{12}$) were used. Z'-factor was calculated using reported methods.[5]

**SI Table 19:** Z'-Factor of TO-PRO-1 for RNA targets

| **Measure** | **HIV-1 TAR** | **HIV-2 TAR** | **$rCAG_{12}$** | **$rCUG_{12}$** |
|---|---|---|---|---|
| Z'-Factor[a]: | 0.92 ± 0.03 | 0.85 ± 0.04 | 0.89 ± 0.04 | 0.85 ± 0.07 |

[a]Values represent the average and standard deviation of three inter-run replicates

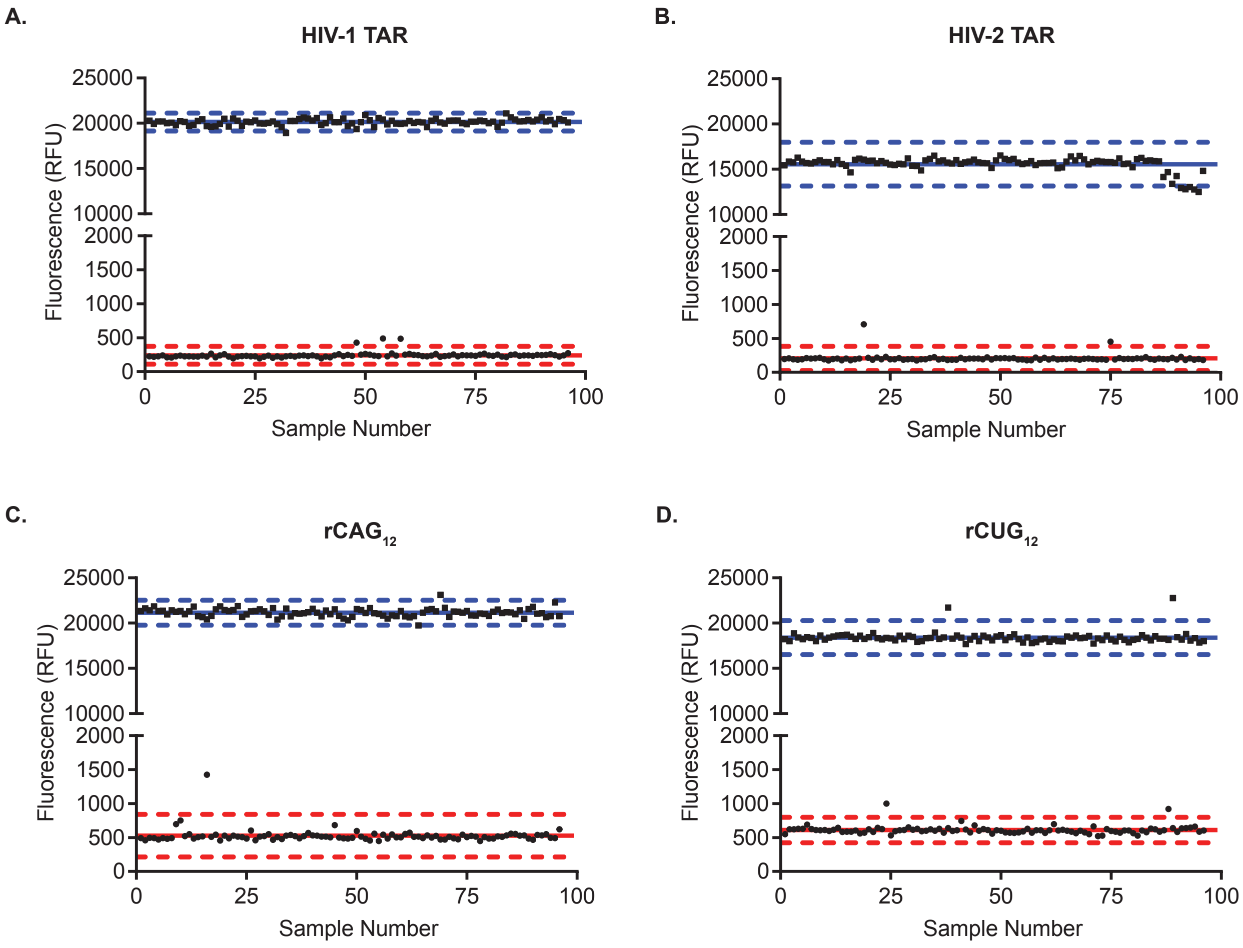

**SI Figure 5.** Representative Z'-factor measurements for TO-PRO-1 indicator displacement assay with HIV-1 TAR (A), HIV-2 TAR (B), $rCAG_{12}$ (C), and $rCUG_{12}$ (D). Fluorescence measurements for 96 positive controls (filled squares) and 96 negative controls (filled circles). All controls contained 500 nM TO-PRO-1, 140 mM KCl, 10 mM $NaH_2PO_4$, 1 mM $MgCl_2$, 0.1 mM EDTA, 0.01% Triton-x-100, and 5% DMSO (pH 7.2). Positive controls contained an RNA concentration of 0.15 μM (HIV-1 TAR), 0.14 μM (HIV-2 TAR), 0.13 μM ($rCAG_{12}$), or 0.18 μM ($rCUG_{12}$). Controls were excited at 492 nm and emission was recorded at 575 nm. Solid lines represent the mean fluorescence for positive and negative controls. Dashed lines represent 3 standard deviations above and below the mean fluorescence.

**Z'-Factor Measurements for Tat Peptide Indicator Displacement Assay**

In a 384-well plate, 96 positive controls (16 µL) were manually plated in alternating columns, containing 50 nM Tat peptide, 140 mM KCl, 10 mM $NaH_2PO_4$, 1 mM $MgCl_2$, 0.1 mM EDTA, 0.01% Triton-x-100, and 5% DMSO (pH 7.2). Similarly, 96 negative controls (16 µL) were manually plated in alternating columns, containing the same reagents and additionally, RNA at a concentration that corresponded to an $Fb^0$ of 0.4: 145 nM (HIV-1 TAR) or 80 nM (HIV-2 TAR). The plate was placed on a shaker at 100 rpm for 2 mins, centrifuged at 4000 rpm for 1 min, and incubated at ambient temperature in the dark for 30 mins. The plate was excited at 485 nm and emission was recorded at 590 nm with 15 nm slits. A read height of 11.5 mm and gain of 2237 (HIV-1 TAR) or 2305 (HIV-2 TAR) were used. Z'-factor was calculated using reported methods.[5]

**SI Table 20:** Z'-Factor of Tat peptide for RNA targets

| Measure | HIV-1 TAR | HIV-2 TAR |
|---|---|---|
| Z'-Factor[a]: | 0.74 ± 0.03 | 0.78 ± 0.06 |

[a]Values represent the average and standard deviation of three inter-run replicates

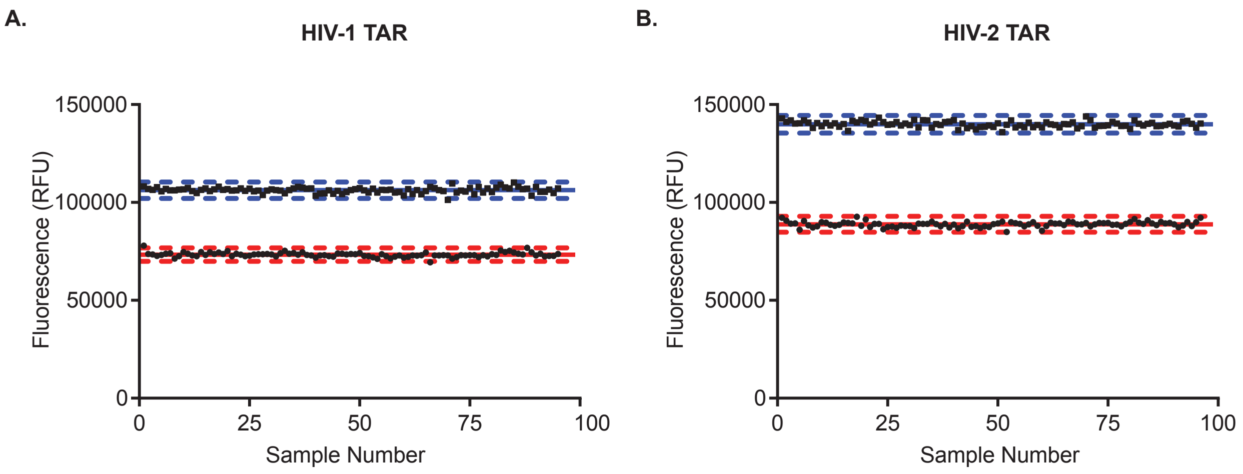

**SI Figure 6.** Representative Z'-factor measurements for Tat peptide indicator displacement assay with HIV-1 TAR (A.) and HIV-2 TAR (B.). Fluorescence measurements for 96 positive controls (filled squares) and 96 negative controls (filled circles). All controls contained 50 nM Tat peptide, 140 mM KCl, 10 mM $NaH_2PO_4$, 1 mM $MgCl_2$, 0.1 mM EDTA, 0.01% Triton-x-100, and 5% DMSO (pH 7.2). Positive controls contained an RNA concentration of 145 nM (HIV-1 TAR) or 80 nM (HIV-2 TAR). Controls were excited at 485 nm and emission was recorded at 590 nm. Solid lines represent the mean fluorescence for positive and negative controls. Dashed lines represent 3 standard deviations above and below the mean fluorescence.

### SI-8. *In Vitro* High-throughput Screening of the DRTL

**Screening of the DRTL with TO-PRO-1 Indicator Displacement Assay**

In a 384-well plate, samples (20 µL) were robotically plated, containing 500 nM TO-PRO-1, 140 mM KCl, 10 mM $NaH_2PO_4$, 1 mM $MgCl_2$, 0.1 mM EDTA, 0.01% Triton-x-100, 5% DMSO, and RNA: 0.15 µM (HIV-1 TAR), 0.14 µM (HIV-2 TAR), 0.13 µM ($rCAG_{12}$), or 0.18 µM ($rCUG_{12}$) (pH 7.2). The plate was centrifuged at 1000 rpm for 1 min and incubated at ambient temperature in the dark for 30 mins. Small molecules (final concentration = 25 µM) were robotically transferred to the 384-well plate. Controls (20 µL) included those that contained only RNA and TO-PRO-1 in buffer (n = 16). The plate was centrifuged at 1000 rpm for 1 min and incubated at ambient temperature in the dark for 30 mins. The plate was excited at 492 nm and emission was recorded at 575 nm with 9 and 15 nm slits, respectively. A read height of 11.3 mm and gain of 1708 (HIV-1 TAR), 1654 (HIV-2 TAR), 1881 ($rCAG_{12}$), or 1964 ($rCUG_{12}$) were used. The percentage of TO-PRO-1 displaced was calculated using Equation (2).

$$Displacement\ (\%) = 100 - \left(100\frac{F_{I+RNA+SM}}{F_{I+RNA}}\right) \quad (2)$$

Where $F_{I+RNA+SM}$ is the fluorescence of samples containing indicator (I), RNA, and small molecule (SM), and $F_{I+RNA}$ is the average fluorescence of samples containing only indicator and RNA.

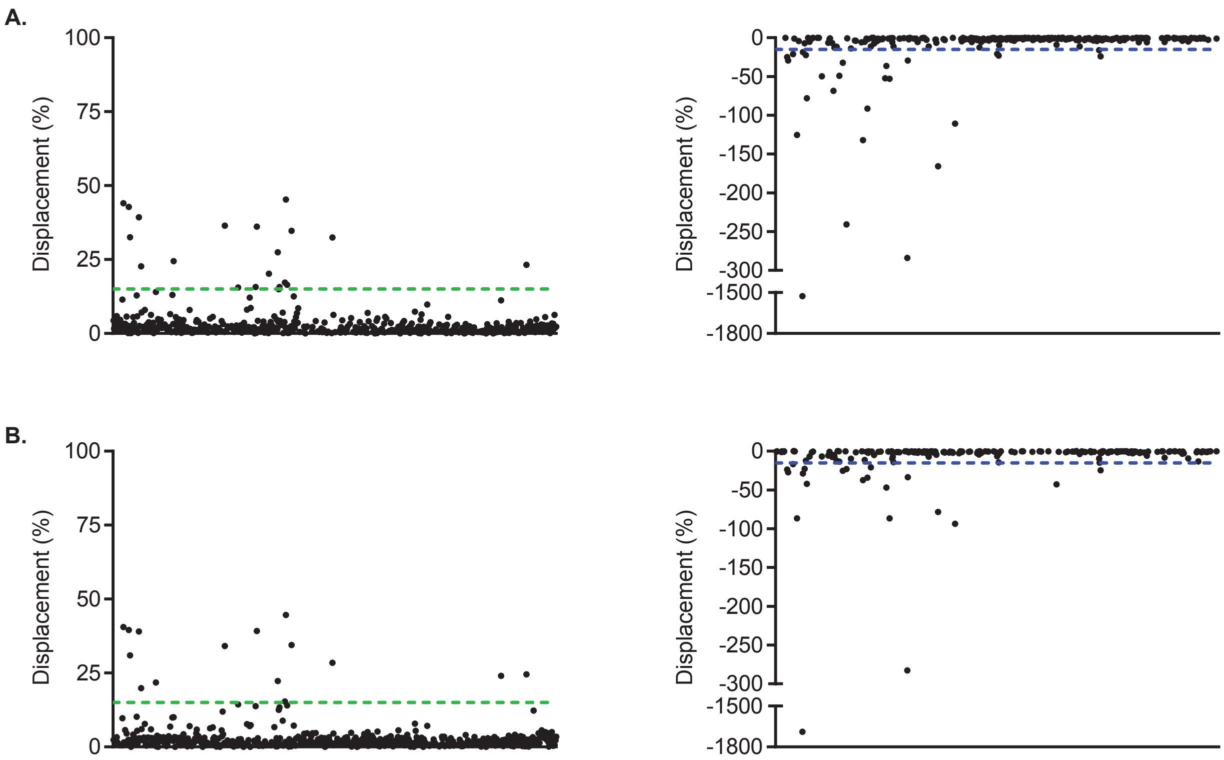

**SI Figure 7.** Percentage of TO-PRO-1 displacement from HIV-1 TAR by the DRTL. The DRTL (black circles) was screened at 25 μM against HIV-1 TAR (0.15 μM) in duplicate (A, B) in the presence of 500 nM TO-PRO-1, 140 mM KCl, 10 mM $NaH_2PO_4$, 1 mM $MgCl_2$, 0.1 mM EDTA, 0.01% Triton-x-100, and 5% DMSO (pH 7.2). Samples were excited at 492 nm and emission was recorded at 575 nm. The percentage of TO-PRO-1 displacement was calculated using equation 1. Green dashed line represents 15% displacement. For ease of visualization, small molecules that induced negative displacement were plotted separately. Small molecules that resulted in < -15% displacement (blue dashed line) were excluded from further consideration.

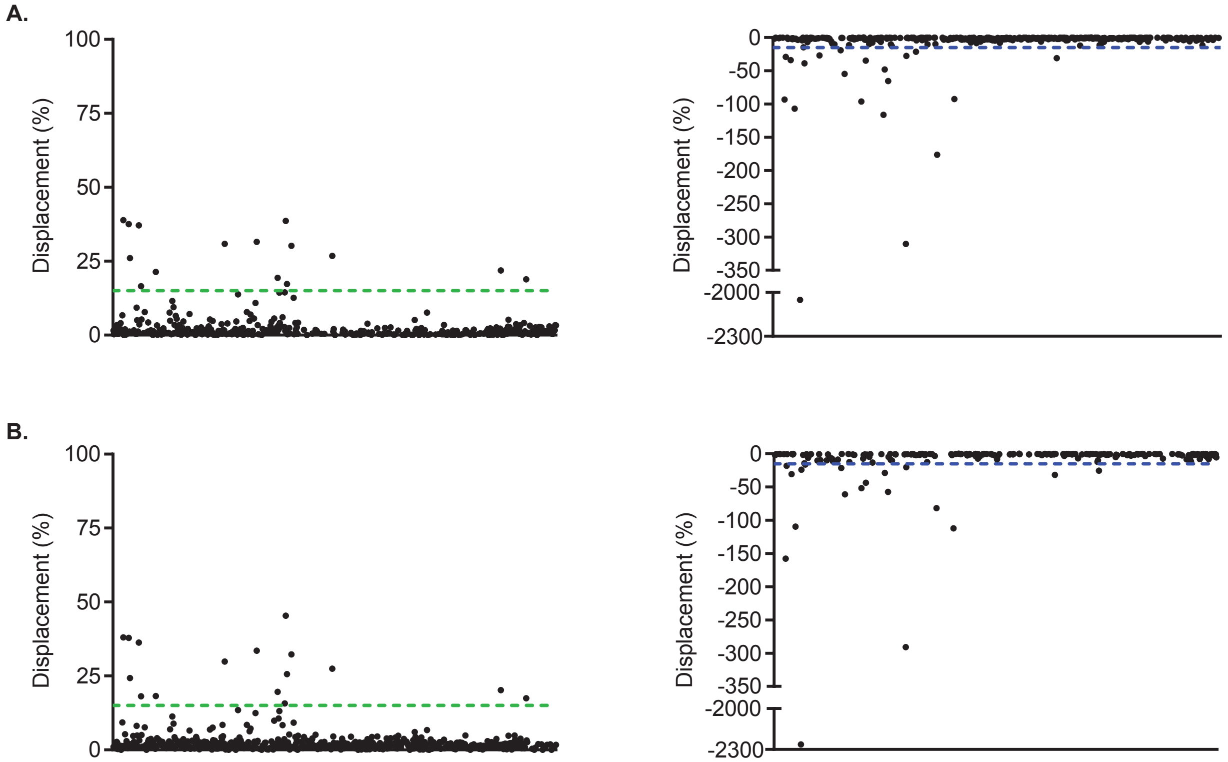

**SI Figure 8.** Percentage of TO-PRO-1 displacement from HIV-2 TAR by the DRTL. The DRTL (black circles) was screened at 25 µM against HIV-2 TAR (0.14 µM) in duplicate (A, B) in the presence of 500 nM TO-PRO-1, 140 mM KCl, 10 mM $NaH_2PO_4$, 1 mM $MgCl_2$, 0.1 mM EDTA, 0.01% Triton-x-100, and 5% DMSO (pH 7.2). Samples were excited at 492 nm and emission was recorded at 575 nm. The percentage of TO-PRO-1 displacement was calculated using equation 1. Green dashed line represents 15% displacement. For ease of visualization, small molecules that induced negative displacement were plotted separately. Small molecules that resulted in < -15% displacement (blue dashed line) were excluded from further consideration.

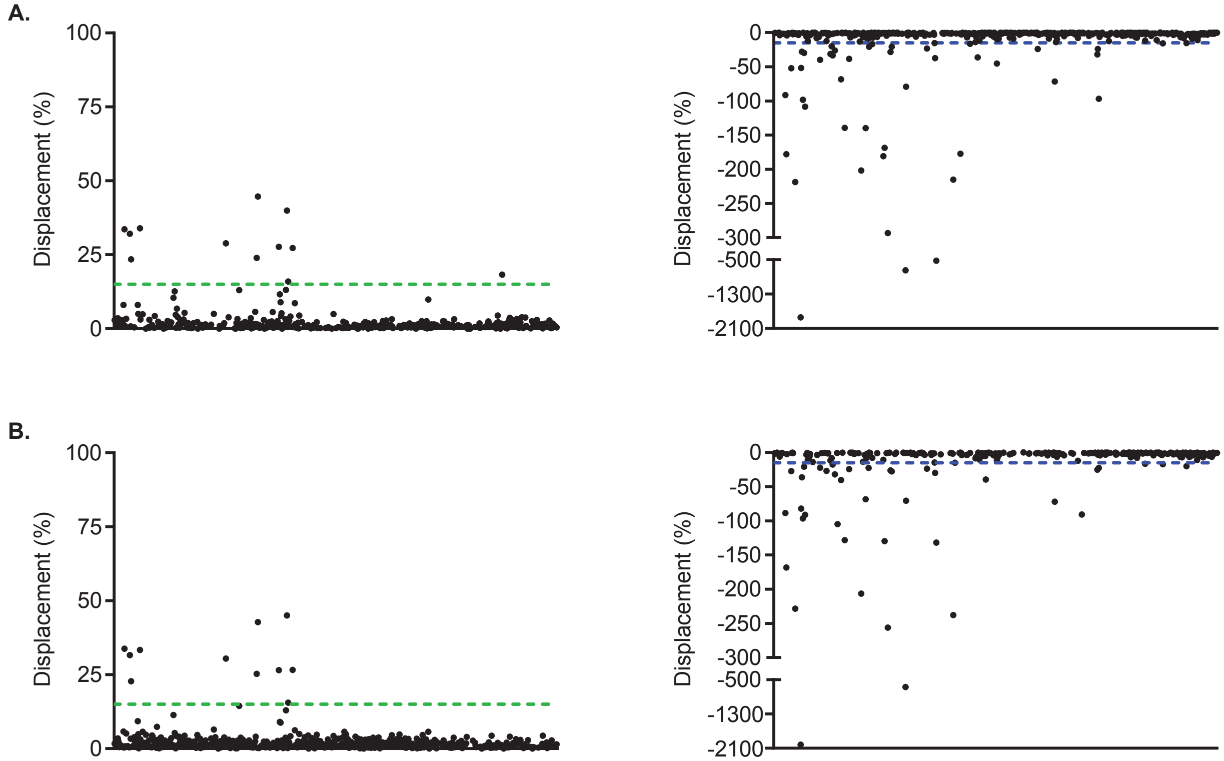

**SI Figure 9.** Percentage of TO-PRO-1 displacement from $rCAG_{12}$ by the DRTL. The DRTL (black circles) was screened at 25 µM against $rCAG_{12}$ (0.13 µM) in duplicate (A, B) in the presence of 500 nM TO-PRO-1, 140 mM KCl, 10 mM $NaH_2PO_4$, 1 mM $MgCl_2$, 0.1 mM EDTA, 0.01% Triton-x-100, and 5% DMSO (pH 7.2). Samples were excited at 492 nm and emission was recorded at 575 nm. The percentage of TO-PRO-1 displacement was calculated using equation 1. Green dashed line represents 15% displacement. For ease of visualization, small molecules that induced negative displacement were plotted separately. Small molecules that resulted in < -15% displacement (blue dashed line) were excluded from further consideration.

A.

B.

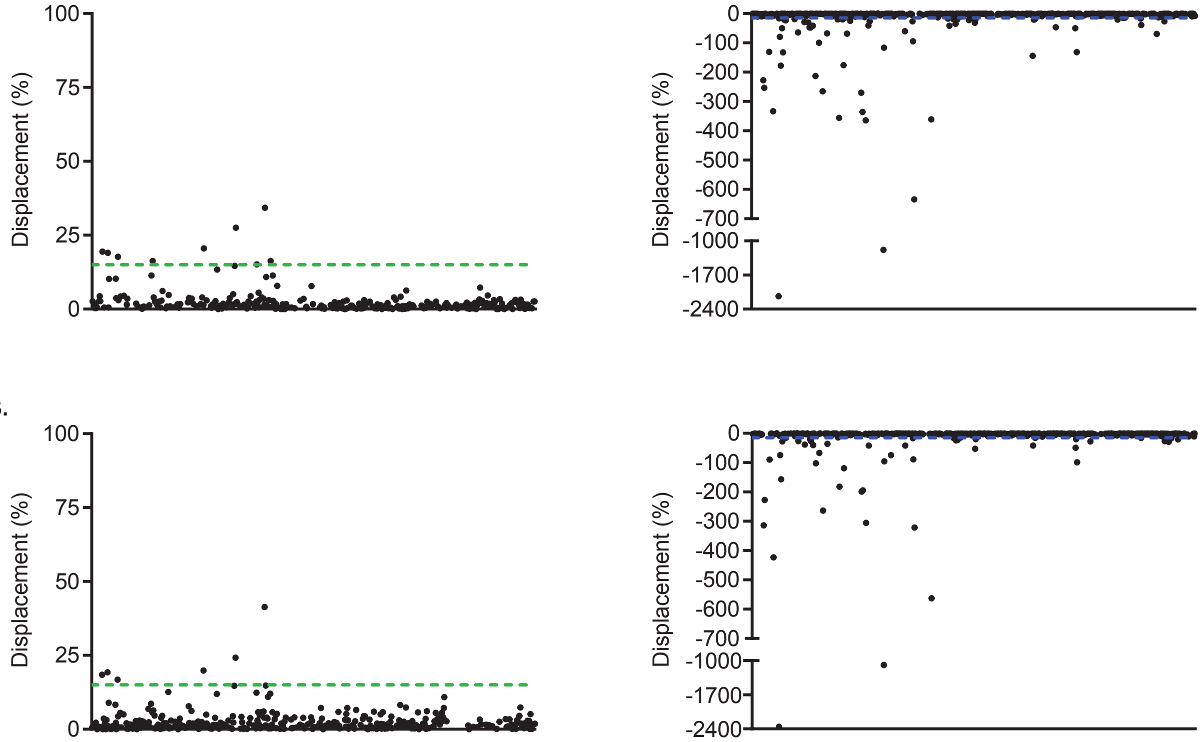

**SI Figure 10.** Percentage of TO-PRO-1 displacement from $rCUG_{12}$ by the DRTL. The DRTL (black circles) was screened at 25 µM against $rCUG_{12}$ (0.18 µM) in duplicate (A, B) in the presence of 500 nM TO-PRO-1, 140 mM KCl, 10 mM $NaH_2PO_4$, 1 mM $MgCl_2$, 0.1 mM EDTA, 0.01% Triton-x-100, and 5% DMSO (pH 7.2). Samples were excited at 492 nm and emission was recorded at 575 nm. The percentage of TO-PRO-1 displacement was calculated using equation 1. Green dashed line represents 15% displacement. For ease of visualization, small molecules that induced negative displacement were plotted separately. Small molecules that resulted in < -15% displacement (blue dashed line) were excluded from further consideration.

**SI Table 21:** Displacement percentages by small molecule hits identified with TO-PRO-1

| | HIV-1 TAR | | HIV-2 TAR | | $rCAG_{12}$ | | $rCUG_{12}$ | |
|---|---|---|---|---|---|---|---|---|
| **Compound** | **Screen 1** | **Screen 2** | **Screen 1** | **Screen 2** | **Screen 1** | **Screen 2** | **Screen 1** | **Screen 2** |
| 1 | 39% | 39% | 37% | 36% | 34% | 33% | 18% | 17% |
| 2 | 36% | 34% | 31% | 30% | 29% | 30% | 21% | 20% |
| 3 | 36% | 39% | 32% | 34% | 45% | 43% | 28% | 24% |
| 4 | 44% | 41% | 39% | 38% | 34% | 34% | 19% | 18% |
| 5 | 43% | 40% | 38% | 38% | 32% | 32% | 19% | 19% |
| 6 | 35% | 34% | 30% | 32% | 27% | 27% | 16% | 12% |
| 7 | 33% | 31% | 26% | 24% | 23% | 23% | 10% | 9% |
| 8 | 32% | 28% | 27% | 27% | 5% | 4% | 8% | 4% |
| 9 | 23% | 25% | 19% | 17% | -3% | -3% | -5% | -1% |
| 10 | 17% | 15% | 14% | 16% | 13% | 13% | 1% | 6% |
| 11 | 16% | 14% | 11% | 12% | 24% | 25% | 15% | 15% |
| 28 | 45% | 45% | 39% | 45% | 40% | 45% | 34% | 41% |
| 29 | 27% | 22% | 19% | 20% | 28% | 27% | 15% | 12% |
| 30 | 16% | 14% | 17% | 26% | 16% | 16% | 11% | 15% |
| 31 | 11% | 24% | 22% | 20% | 18% | -4% | 7% | 3% |

**Screening of the DRTL with Tat Peptide Indicator Displacement Assay**

In a 384-well plate, samples (20 µL) were robotically plated, containing 50 nM Tat peptide, 140 mM KCl, 10 mM $NaH_2PO_4$, 1 mM $MgCl_2$, 0.1 mM EDTA, 0.01% Triton-x-100, 5% DMSO, and RNA: 145 nM (HIV-1 TAR) or 80 nM (HIV-2 TAR) (pH 7.2). The plate was centrifuged at 1000 rpm for 1 min and incubated at ambient temperature in the dark for 30 mins. Small molecules (final concentration = 25 µM) were robotically transferred to the 384-well plate. Controls (20 µL) included those that contained only RNA and Tat peptide in buffer (n = 16). The plate was centrifuged at 1000 rpm for 1 min and incubated at ambient temperature in the dark for 30 mins. The plate was excited at 485 nm and emission was recorded at 590 nm with 15 nm slits. A read height of 11.3 mm and gain of 2237 (HIV-1 TAR) or 2305 (HIV-2 TAR) were used. The percentage of Tat peptide displaced was calculated using Equation (3).

$$Displacement\ (\%) = 100 - \left(100\,\frac{F_{I+RNA+SM} - F_I}{F_{I+RNA} - F_I}\right) \quad (3)$$

Where $F_{I+RNA+SM}$ is the fluorescence of samples containing indicator (I), RNA, and small molecule (SM), $F_I$ is the average fluorescence of samples containing only indicator, and $F_{I+RNA}$ is the average fluorescence of samples containing only indicator and RNA.

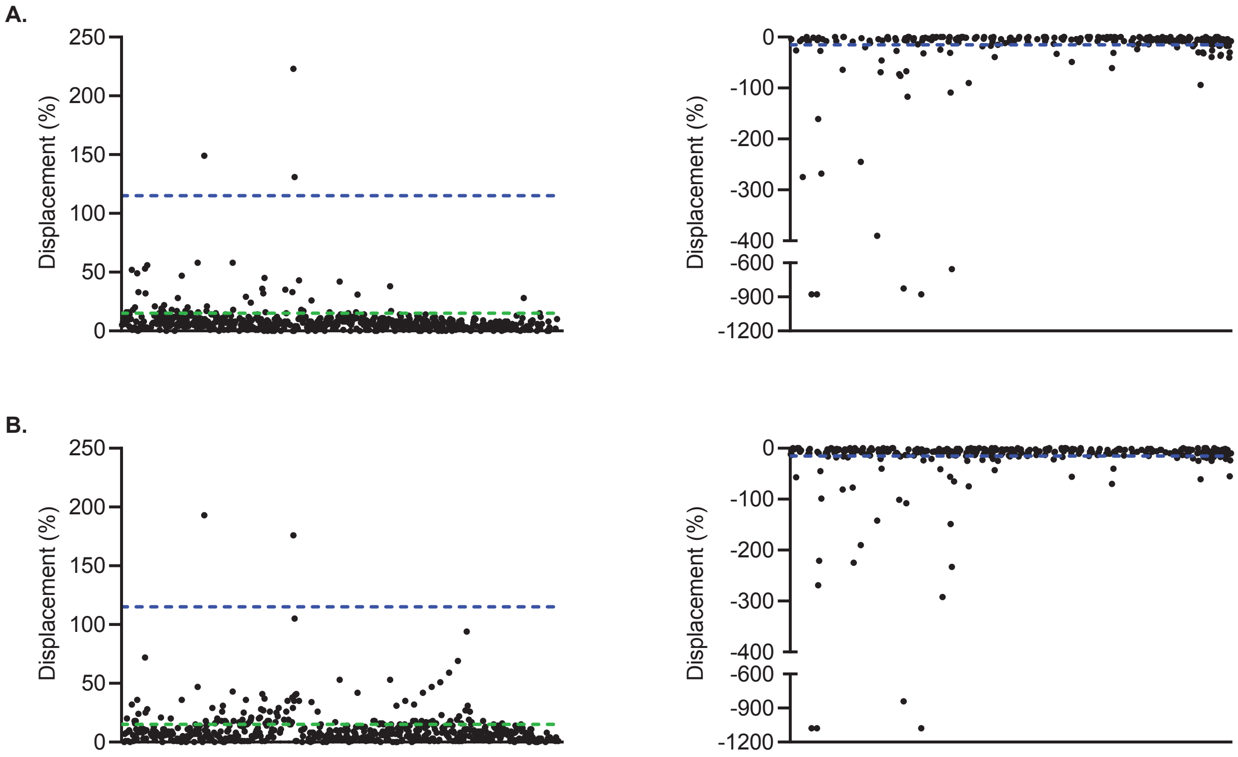

**SI Figure 11.** Percentage of Tat peptide displaced from HIV-1 TAR by the DRTL. The DRTL (black circles) was screened at 25 μM against HIV-1 TAR (145 nM) in duplicate (A., B.) in the presence of 50 nM Tat peptide, 140 mM KCl, 10 mM $NaH_2PO_4$, 1 mM $MgCl_2$, 0.1 mM EDTA, 0.01% Triton-x-100, and 5% DMSO (pH 7.2). Samples were excited at 485 nm and emission was recorded at 590 nm. The percentage of Tat peptide displacement was calculated using equation 2. Green dashed line represents 15% displacement. For ease of visualization, small molecules that induced negative displacement were plotted separately. Small molecules that resulted in either > 115% or < -15% displacement (blue dashed lines) were excluded from further consideration.

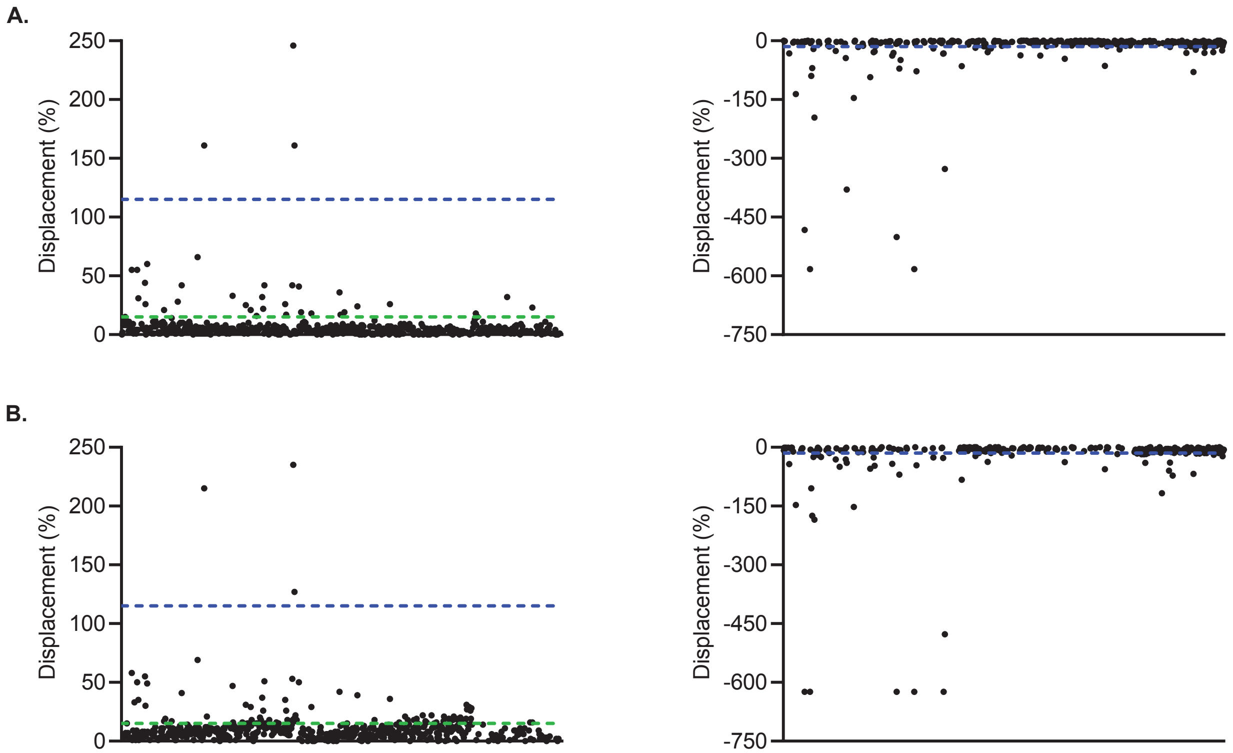

**SI Figure 12.** Percentage of Tat peptide displaced from HIV-2 TAR by the DRTL. The DRTL (black circles) was screened at 25 μM against HIV-2 TAR (80 nM) in duplicate (a, b) in the presence of 50 nM Tat peptide, 140 mM KCl, 10 mM $NaH_2PO_4$, 1 mM $MgCl_2$, 0.1 mM EDTA, 0.01% Triton-x-100, and 5% DMSO (pH 7.2). Samples were excited at 485 nm and emission was recorded at 590 nm. The percentage of Tat peptide displacement was calculated using equation 2. Green dashed line represents 15% displacement. For ease of visualization, small molecules that induced negative displacement were plotted separately. Small molecules that resulted in either > 115% or < -15% displacement (blue dashed lines) were excluded from further consideration.

**SI Table 22:** Displacement percentages by small molecule hits identified with Tat peptide

| | HIV-1 TAR | | HIV-2 TAR | |
|---|---|---|---|---|
| **Compound** | **Screen 1** | **Screen 2** | **Screen 1** | **Screen 2** |
| 1 | 56% | 28% | 60% | 49% |
| 2 | 58% | 43% | 33% | 47% |
| 3 | 45% | 37% | 42% | 51% |
| 4 | 52% | 32% | 55% | 58% |
| 5 | 49% | 36% | 55% | 50% |
| 6 | 43% | 35% | 41% | 50% |
| 7 | 33% | 24% | 31% | 35% |
| 8 | 42% | 53% | 36% | 42% |
| 9 | 9% | 4% | 23% | 16% |
| 10 | 33% | 38% | 42% | 53% |
| 11 | 32% | 27% | 22% | 26% |
| 12 | 47% | 36% | 42% | 41% |
| 13 | 29% | 36% | 25% | 31% |
| 14 | 15% | 26% | 17% | 26% |
| 15 | 38% | 53% | 26% | 36% |
| 16 | 31% | 42% | 24% | 39% |
| 17 | 26% | 34% | 18% | 29% |
| 18 | 53% | 72% | 44% | 55% |
| 19 | 58% | 47% | 66% | 69% |
| 20 | 32% | 25% | 26% | 30% |
| 21 | 24% | 19% | 21% | 29% |
| 22 | 16% | 16% | 6% | 15% |
| 23 | 17% | 15% | 7% | 3% |
| 24 | 20% | 16% | 4% | 6% |
| 25 | 36% | 41% | 32% | 37% |
| 26 | 22% | 11% | 21% | 17% |
| 27 | 21% | 15% | 10% | 21% |

**SI Table 23:** R-BIND (SM) ligands that were nearest neighbors to small molecule hits

| R-BIND (SM) | Target | Hit(s) |
|---|---|---|
| 0013 | yjdF Riboswitch | 8, 25 |
| 0016 | A Disintegrin and Metalloproteinase 10 (ADAM10) G-Quadruplex Forming Sequence | 3, 5-7, 11, 22, 29 |
| 0017 | A Disintegrin and Metalloproteinase 10 (ADAM10) G-Quadruplex Forming Sequence | 1, 4-7, 9, 29 |
| 0018 | A Disintegrin and Metalloproteinase 10 (ADAM10) G-Quadruplex Forming Sequence | 13 |
| 0019 | A Disintegrin and Metalloproteinase 10 (ADAM10) G-Quadruplex Forming Sequence | 13 |
| 0021 | DDPAC Microtubule-Associated Protein Tau (MAPT) pre-mRNA | 17, 28, 30 |
| 0028 | Dystrophia Myotonica Protein Kinase (DMPK) r(CUG) repeats | 18 |
| 0029 | Dystrophia Myotonica Protein Kinase (DMPK) r(CUG) repeats | 26 |
| 0030 | Dystrophia Myotonica Protein Kinase (DMPK) r(CUG) repeats | 12, 15, 27 |
| 0031 | Dystrophia Myotonica Protein Kinase (DMPK) r(CUG) repeats | 19 |
| 0033 | Dystrophia Myotonica Protein Kinase (DMPK) r(CUG) repeats | 16 |
| 0034 | Dystrophia Myotonica Protein Kinase (DMPK) r(CUG) repeats | 28, 30 |
| 0035 | Dystrophia Myotonica Protein Kinase (DMPK) r(CUG) repeats | 28, 30 |
| 0036 | Serotonin 2C Receptor (HTR2C) pre-mRNA | 1, 2, 4-7, 9, 14, 24, 29, 31 |
| 0049 | HIV-1 Trans-Activation Response Element (TAR) | 27 |
| 0050 | HIV-1 Trans-Activation Response Element (TAR) | 21 |
| 0052 | HIV-1 Trans-Activation Response Element (TAR) | 20, 21 |
| 0053 | HIV-1 Trans-Activation Response Element (TAR) | 21 |
| 0056 | HIV-1 Trans-Activation Response Element (TAR) | 1, 2 |
| 0060 | HIV-1 Trans-Activation Response Element (TAR) | 27, 28, 30 |
| 0066 | Japanese Encephalitis Virus (JEV) Frameshift Site | 10, 23 |

### SI-9. References

(1) Wenderski, T. A., Stratton, C. F., Bauer, R. A., Kopp, F., and Tan, D. S. (2015) Principal component analysis as a tool for library design: a case study investigating natural products, brand-name drugs, natural product-like libraries, and drug-like libraries. *Methods Mol. Biol. 1263*, 225-242.

(2) Mitchell, J. B. (2014) Machine learning methods in chemoinformatics. *Wiley Interdiscip. Rev. Comput. Mol. Sci. 4*, 468-481.

(3) Morgan, B. S., Sanaba, B. G., Donlic, A., Karloff, D. B., Forte, J. E., Zhang, Y., and Hargrove, A. E. (2019) R-BIND: An Interactive Database for Exploring and Developing RNA-Targeted Chemical Probes. *ACS Chem. Biol. 14*, 2691-2700.

(4) Del Villar-Guerra, R., Gray, R. D., Trent, J. O., and Chaires, J. B. (2018) A rapid fluorescent indicator displacement assay and principal component/cluster data analysis for determination of ligand-nucleic acid structural selectivity. *Nucleic Acids Res. 46*, e41.

(5) Zhang, J. H., Chung, T. D., and Oldenburg, K. R. (1999) A Simple Statistical Parameter for Use in Evaluation and Validation of High Throughput Screening Assays. *J. Biomol. Screen. 4*, 67-73.
